# Multi-omics analysis reveals how the natural compound bostrycin induces iron starvation and ferroptosis

**DOI:** 10.64898/2026.11.24.768891

**Authors:** Lina-Marie Wagner, Azmal Syed Ali, Isabelle Becher, Julie Haenlin, Adriano de Britto Chaves Filho, Mikhail M. Savitski, Almut Schulze, Jeroen Krijgsveld

## Abstract

The development of novel anticancer drugs remains urgent due to limited efficacy of existing therapies. Natural products are a rich source for discovery of anticancer agents, yet their mechanisms often remain elusive. Bostrycin, a fungal anthraquinone with broad cytotoxicity, is one such compound. Here, we combine global and nascent proteomics with metabolomics to elucidate the mechanism of bostrycin-induced cell death. We show that bostrycin induces ferroptosis, and use thermal proteome profiling to identify thioredoxin reductase 1 (TXNRD1), a key enzyme in glutathione metabolism and redox control, as bostrycin’s primary direct target. Bostrycin triggered an iron-starvation response and reduced iron–sulfur cluster–containing proteins, consistent with impaired cysteine availability. While the resulting iron overload likely triggered ferroptotic death, this coincided with broader disruption of redox balance, glutathione metabolism, and energy pathways. These findings establish bostrycin’s mechanism of action and identify it as a ferroptosis-inducing agent acting distinctly from canonical inducers.

**Highlights:**

- Nascent proteomics across eight cell lines shows that bostrycin induces ferroptosis
- 2D thermal proteome profiling identifies TXNRD1 as the direct target of bostrycin
- Bostrycin depletes cysteine and glutathione and reduces Fe-S cluster proteins
- Bostrycin triggers an iron-starvation response not seen with IKE or RSL3

## Introduction

Despite major advances in cancer therapy, the development of novel anticancer drugs remains essential. This need arises from cancer types for which effective treatments are still unavailable, the considerable heterogeneity within and between tumors that complicates personalized treatment, and the frequent emergence of resistance that limits the long-term efficacy of existing therapies. As a result, combination therapies are increasingly being pursued, leveraging synergistic interactions between complementary agents to enhance therapeutic efficacy and improve patient outcomes ^1^.

Natural products constitute a historically important source of anticancer agents. Derived from plants, animals, and microorganisms, they exhibit extensive chemical diversity and have served as templates for many highly successful (semi-)synthetic drugs ^2, 3, 4, 5^. Prominent examples include paclitaxel, camptothecin, cytarabine, and their derivatives, which are widely used across multiple cancer indications, with many others currently under clinical evaluation ^6, 7^. The promise of natural products lies in their capacity to enable innovative therapeutic interventions by targeting cellular pathways that may not be rationally anticipated; however, this same feature often renders mechanistic elucidation particularly challenging. As a result, mechanistic dissection is typically reserved for compounds that show robust and reproducible effects across multiple model systems and phenotypic readouts ^6^.

One such compound is bostrycin, a pigmented anthraquinone metabolite isolated from several fungal species. Initially characterized for its chemical structure ^8^ and antibiotic and phytotoxic activities ^9, 10^, bostrycin was shown to trigger apoptosis in yeast, induce cell cycle arrest, increase reactive oxygen species (ROS), and disrupt mitochondrial function ^11^. Subsequent studies demonstrated that bostrycin inhibits proliferation and induces cell death in a range of human cancer cell lines ^12, 13, 14^. However, mechanistic interpretations have remained inconsistent. Several reports implicated PI3K/AKT signaling, however contradictorily showing a decrease ^13^ and increase of pAKT ^12^. A zebrafish embryo screen further suggested that bostrycin inhibits PI3K/AKT based on the observation that bostrycin and the established PI3K/AKT inhibitor LY294002 similarly reduced body axis extension ^15^. In the bacterial pathogen *Mycobacterium tuberculosis*, proteomic analyses linked bostrycin to oxidative stress and metabolic disruption, and molecular docking simulations suggested Rv3684, a cysteine desulfhydrase, and Rv1908c, a bifunctional catalase and peroxidase, as the most likely direct targets of bostrycin ^16^. As these proteins lack human orthologues, the molecular targets of bostrycin in human cells remain unresolved.

Collectively, these studies established bostrycin as a cytotoxic compound with potential for anticancer development, yet its mechanism of action in human cells has remained unclear. Here, we address these gaps using an integrated multi-omics approach combining global and nascent proteomics with thermal proteome profiling, metabolomics and functional assays. We show that bostrycin induces a strong iron starvation response and ferroptosis across multiple human cancer cell lines and used thermal proteome profiling to identify thioredoxin reductase 1 (TXNRD1) as its primary target. Functional validation confirmed key features of ferroptosis, including lipid peroxidation and membrane blebbing, while omics analyses revealed coordinated changes in glutathione metabolism, redox regulation, energy metabolism, and iron homeostasis. Together, these findings demonstrate the power of integrated proteomic and metabolomic strategies to resolve the mechanisms of previously uncharacterized compounds, including ferroptosis, a pathway notable for its mechanistic complexity and cell type–dependence ^17^. Given the growing recognition of ferroptosis as a therapeutically exploitable vulnerability in cancer ^18^, our work provides important insight into bostrycin as a lead compound capable of inducing ferroptosis-mediated cell death.

## Results

### Bostrycin is cytotoxic for breast cancer cell lines but without consistent proteome alteration

To investigate how bostrycin (**Figure 1A**) induces cell death, we selected a panel of eight breast cancer cell lines to conduct a series of molecular and functional assays to determine bostrycin-induced cellular effects. Cell viability assays revealed IC50 values within a narrow range of 5 - 12 µM (**Figure 1B**), indicating relatively high potency for a non-optimized natural product based on established activity classifications ^19^.

**Figure 1.**
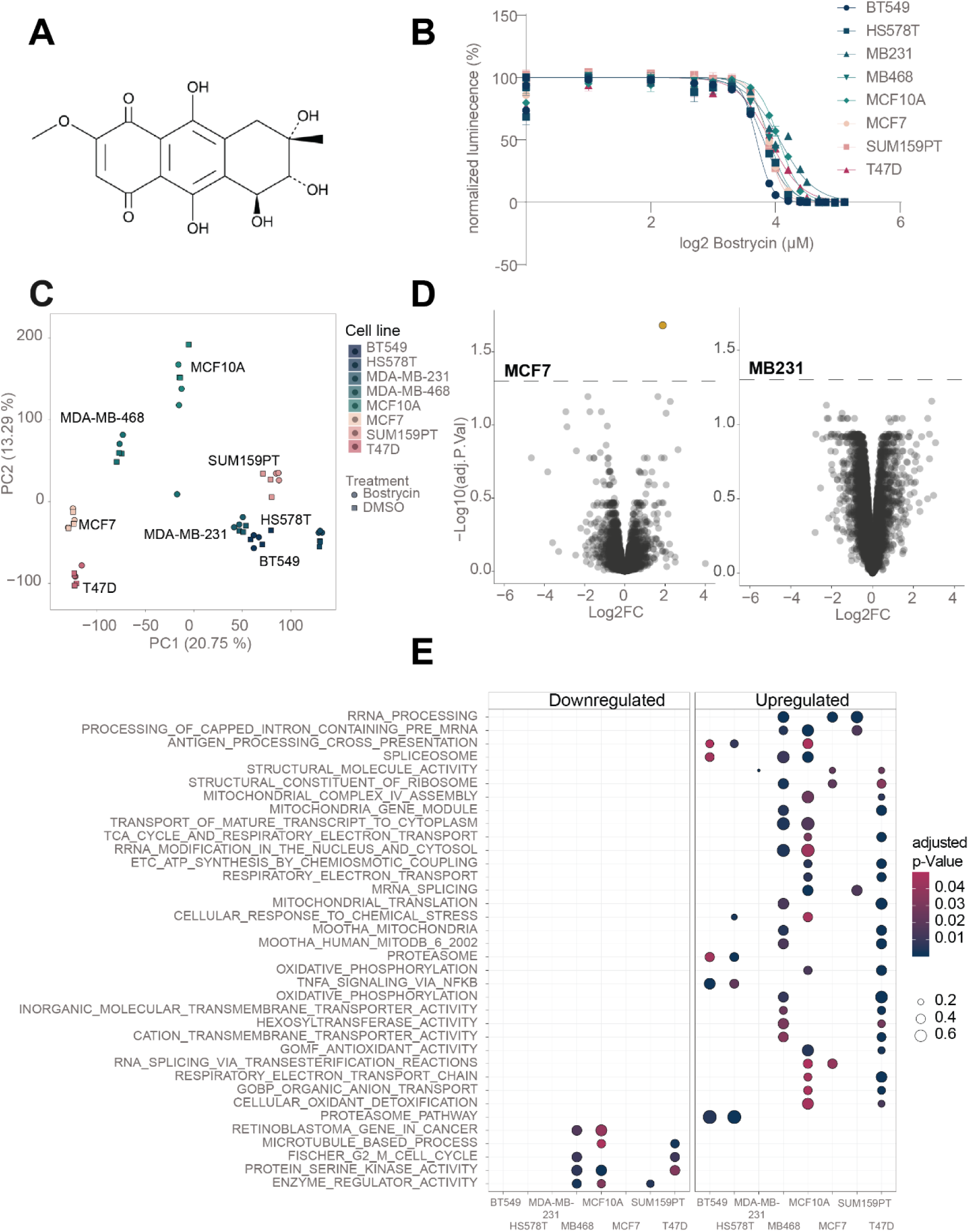
Cell viability and proteome effects of cancer cells treated with bostrycin. **A.** Structure of bostrycin. **B.** Dose-response curves of cell viability after treating with different concentrations of the natural compound bostrycin for 72h. **C.** Principal Component Analysis (PCA) of proteomic data from eight cell lines treated with bostrycin. **D.** Volcano plots indicating differential protein expression in MCF7 (left) and MDA-MB-231 cells (right) after treatment with bostrycin. Highlighted are proteins that are significantly differentially expressed after multiple hypothesis testing. **E.** Gene set enrichment analysis of proteomic data in eight cell lines after treatment with bostrycin, across the MSigDB databases Hallmark, C2 and C5. Shown are terms that were enriched in at least 2 cell lines.

To identify cellular processes underlying bostrycin-induced cell death, we performed proteome profiling of the eight cell lines upon bostrycin treatment compared to DMSO-treated controls. Cells were treated in triplicate with bostrycin at their respective IC₅₀ concentrations for 6 h and analyzed by label-free proteomics. In total, more than 10,000 protein groups were identified across all cell lines, with ∼7,000–8,000 proteins detected per line. HS578T cells consistently yielded the lowest coverage (6,835 proteins), whereas T47D cells showed the highest (8,332; **Supplementary Figure S1; Table 1**), likely reflecting cell type–intrinsic proteome characteristics.

Principal component analysis (PCA) indicated that cell type, rather than treatment, was the dominant source of variance, as expected for a short treatment duration (**Figure 1C**). To assess bostrycin-induced effects within individual cell lines, we performed differential protein expression analyses. This showed that very few proteins (e.g. in MCF7 cells) or no proteins at all (e.g. in MDA-MB-231 cells) reached statistical significance (adjusted p-value <0.05) **(Figure 1D)**. This trend extended across the remaining cell lines (**Supplementary Figure S1B)**. Using a less stringent cutoff (>2-fold change with a nominal p value <0.05 prior to multiple testing correction), 100–200 dysregulated proteins per cell line were identified (**Supplementary Table S1**). However, the overlap of commonly up- or downregulated proteins across cell lines was minimal (**Supplementary Figures S1C and D**). Gene set enrichment analysis (GSEA) of ranked proteomic data from each cell line yielded a diverse set of enriched terms (**Figure 1E**), including categories related to electron transport and RNA metabolism; however, none were shared by more than three cell lines. Together, these results indicate that bostrycin elicits modest and cell-line-specific proteomic alterations, precluding the identification of a consistent global response.

### Nascent proteome analysis indicates that bostrycin induces ferroptosis

As an alternative strategy to characterize the proteomic response to bostrycin, we performed a nascent proteome analysis. This approach focuses on newly synthesized proteins following a cellular perturbation and can reveal early proteomic changes that may not yet be detectable at the steady-state level ^20, 21^. To this end, all eight cell lines were treated with bostrycin for 6 and 12 hours, and were pulse-labeled with azido-homoalanine (AHA) and SILAC amino acids during the last 6 hours of treatment. This was done analogously for non-treated controls (**Figure 2A**). Following click-chemistry enrichment of nascent proteins and mass spectrometric analysis, we quantified 2,000– 3,000 proteins per cell line (approximately 6,500 in total) based on their SILAC ratios (**Supplementary Figure S2A; Supplementary Table S2**). PCA revealed that samples clustered primarily by cell line; however, several lines, including MDA-MB-231, MCF10A, and SUM159PT, showed partial separation between the 6-hour and 12-hour treatments (**Figure 2B**), indicating a detectable treatment-induced proteomic effect.

**Figure 2.**
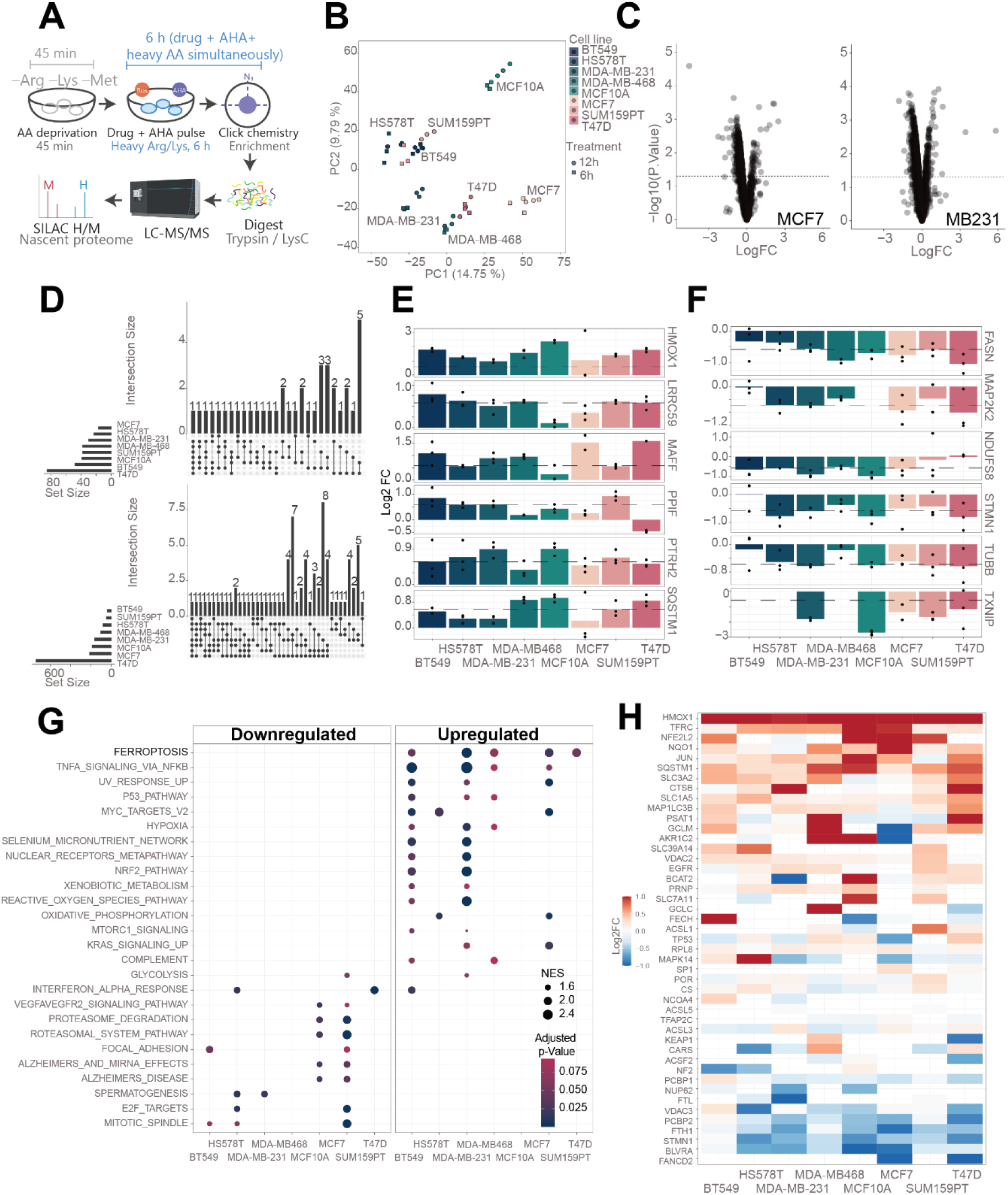
Nascent proteomics of bostrycin-treated cancer cell lines. **A.** Experimental scheme. **B.** Principal Component Analysis (PCA) of nascent proteomic data from eight cell lines treated with bostrycin for 6h or 12h. **C.** Volcano plots indicating differential protein expression in nascent proteomic data of bostrycin-treated MCF7 (left) and MDA-MB-231 cells (right). **D.** Upset plots indicating the overlap in upregulated (left) and downregulated proteins (right) in the nascent proteome of bostrycin-treated cancer cell lines. **E, F.** Selection of the most common up (E) and down-regulated (F) proteins in the nascent proteome of bostrycin-treated cancer cell lines. **G.** Gene set enrichment analysis of newly synthesized proteins after 12h of bostrycin treatment. **H.** Tile plot of proteins that are associated with ferroptosis. Shown are the average log2-fold changes from triplicate nascent proteome analysis of bostrycin-treated cell lines.

Limma analysis was used to identify significantly altered proteins (**Figure 2C**), applying a >2-fold change and adjusted p < 0.05 cutoff. Across the cell lines and for both time points, more proteins were downregulated (6h: 20-140; 12h: 40-740) than upregulated (6h: 10-30; 12h: 20-80), and more de-regulated proteins were observed at 12h than at 6h (**Supplementary Figure S2B-E**), indicating a progressively stronger response.

In contrast to the global proteome analysis (**Supplementary Figure S1D, E**), several differentially regulated proteins were shared across up to seven cell lines **(Figure 2D)**. Notable upregulated proteins included MAFF, SQSTM1, and HMOX1, all three of which operate in the Nrf2 pathway activated under oxidative stress conditions ^22 23^ (**Figure 2E**). TXNIP, a thioredoxin-interacting protein and oxidative stress mediator, was commonly downregulated (**Figure 2F**). Together, these findings implicate oxidative stress as a key response to bostrycin.

To explore this further, we performed GSEA on the 12 h nascent proteome for each of the eight cell lines. Downregulated proteins did not yield any terms enriched in more than two lines (**Figure 2G**). Among upregulated proteins, the top-scoring term—WP_Ferroptosis—was enriched in five cell lines, the highest number of cell lines sharing any single enriched term in this analysis, suggesting that bostrycin induces cell death via ferroptosis, an iron-dependent mechanism of regulated cell death. To seek additional support, we compiled proteins from the Wikipathway “Ferroptosis” term, supplemented with ferroptosis-associated proteins reported in FerrDb ^24^, and mapped them onto our nascent proteome. This revealed sets of both up- and down-regulated proteins with reasonable consistency across cell lines (**Figure 2H**). The five strongest upregulated proteins—HMOX1, TFRC, NRF2, NQO1, and JUN—are all regulators or effectors of oxidative or iron stress responses. The most downregulated proteins included FTH (ferritin heavy chain) and PCBP2, both involved in iron metabolism, the latter serving as a cytosolic iron chaperone that delivers iron to ferritin for storage ^25^. In addition, downregulation of FANCD2 has been previously associated with induction of ferroptosis ^26^, BLVRA is involved in heme metabolism like HMOX1, and STMN1 has been reported to inhibit ferroptosis in hepatocellular carcinoma ^27^. These data strongly indicate that bostrycin induces an oxidative stress response that activates NRF2, a master regulator that orchestrates redox balance and iron homeostasis, leading to ferroptosis when this balance is disrupted ^28, 29^. Strikingly, treatment of MCF7 cells with the known ferroptosis inducers IKE and RSL3 produced highly distinct nascent proteome responses compared to bostrycin as evidenced by their clear separation in a PCA analysis (**Supplementary Figure S3A; Supplementary Table S3)** and by the very limited overlap in the proteins (**Supplementary Figure S3B)** and cellular processes they regulate (**Supplementary Figure S3C**). Collectively this suggests that bostrycin activates NRF2 and induces ferroptosis, however in a manner that differs from the canonical ferroptosis inducers IKE and RSL3. Although NRF2 activation is generally cytoprotective, the notion that bostrycin induces cell death indicates that the oxidative and ferroptotic insult exceeds NRF2’s protective capacity.

### Bostrycin induces functional hallmarks of ferroptosis

We next performed functional and morphological assays to confirm that bostrycin induces ferroptosis, using MCF7 and MDA-MB-231 cells as model systems. Prolonged bostrycin treatment caused prominent macro-blebbing in both cell lines (**Figure 3A**), a characteristic feature of ferroptotic cell death previously reported in MDA-MB-231 cells ^30^ and distinct from apoptotic blebbing ^30, 31^. Consistently, macro-blebbing was also observed upon treatment with IKE, an inhibitor of system xc⁻ and established ferroptosis inducer ^32^ (**Figure 3A**).

**Figure 3.**
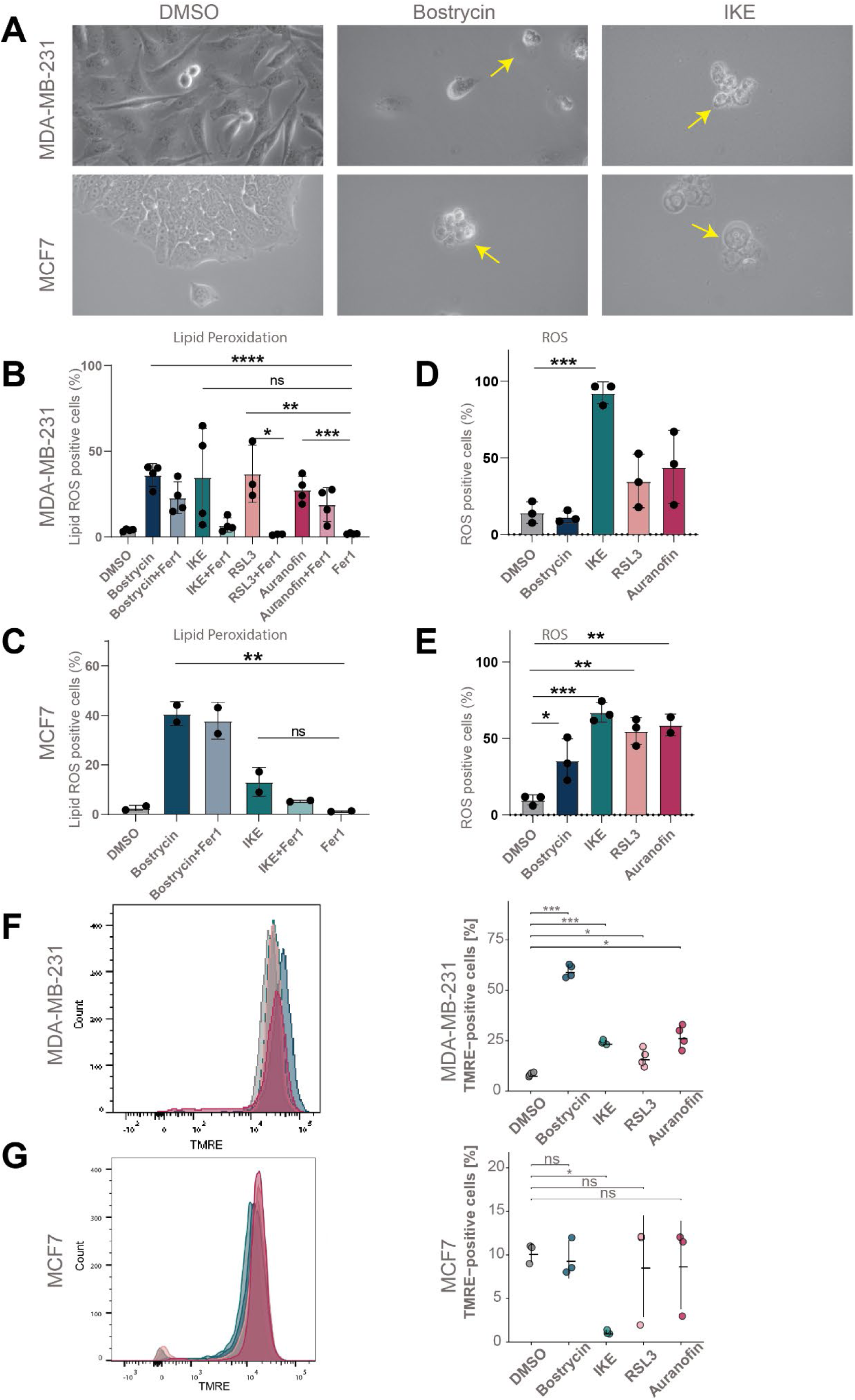
Functional analysis of bostrycin-treated cells. **A.** Light microscopy images of MDA-MB-231 (top row) and MCF7 (bottom row) cells treated with DMSO, bostrycin (50 µM and 20 µM, respectively) for 72h, or with IKE (2.5 µM and 60 µM, respectively) for 27h. Arrows indicate cells that exhibit macroblebbing. **B, C.** Bar plots indicating lipid peroxidation in MDA-MB-231 cells (B) and MCF7 cells (C), assessed as the ratio of lROS positive cells compared to all detected single cells. Shown are treatments with bostrycin (2xIC50), IKE (IC50), RSL3 (2xIC50) and Auranofin (IC50) and co-treatments of the drugs with Fer-1 (20 µM), all for 48h. Error bars indicate the standard deviation of four independent replicates. **D, E.** Bar plots indicating ROS production in MDA-MB-231 cells (D) and MCF7 cells (E), assessed as the ratio of ROS positive cells compared to all detected single cells. **F.** Histogram (left) and bar plot (right) indicating mitochondrial membrane potential measurement using TMRE and flow cytometry in MDA-MB-231 cells upon treatment with bostrycin (2xIC50) IKE (IC50), RSL3 (2xIC50) and Auranofin (IC50). **G.** As in F, for MCF7 cells.

A key hallmark of ferroptosis is the increase in lipid peroxidation that, if not rescued, leads to membrane rupture and cell death ^17^. Since lipid peroxidation can be accompanied by an increase in general ROS, we assessed both ROS and lipid peroxidation using 2’,7’-Dichlordihydrofluorescein-diacetat (H2DCFDA) and C11-Bodipy as probes, respectively, both in MDA-MB-231 and MCF7 cells. Treatment of MDA-MB-231 cells with bostrycin caused a robust increase in lipid peroxidation, comparable to or exceeding that induced by the established ferroptosis inducers IKE, RSL3, and auranofin (**Figure 3B**). This effect was fully rescued by the ferroptosis inhibitor Fer-1 in cells treated with IKE and RSL3, while partial rescue was observed for Bostrycin and Auranofin (**Figure 3B**). The incomplete rescue likely reflects the broader redox disruption caused by these agents. In MCF7 cells, bostrycin induced much stronger lipid peroxidation than IKE (**Figure 3C**), collectively showing that bostrycin elicits a key ferroptotic hallmark in both cell lines. A difference between the cell lines was observed in the partial rescue by Fer-1 in MDA-MB-231 cells that was further diminished in MCF7 cells (**Figure 3B, C and D**). Differences between cell lines were also observed in ROS production: MDA-MB-231 cells responded strongly to IKE, less to RSL3 and auranofin, and not to bostrycin (**Figure 3D**), while MCF7 cells showed a partial response to all four treatments, including bostrycin (**Figure 3E**).

We next assessed treatment effects on hyperpolarization of mitochondrial membrane potential (MMP), which has been associated with induction of ferroptosis ^33^. MDA-MB-231 cells showed increased MMP upon treatment with all four ferroptosis inducers, with the strongest effect observed for bostrycin (**Figure 3F**). In contrast, none of the treatments significantly altered the MMP in MCF7 cells (**Figure 3G**).

Collectively, these experiments show that bostrycin elicits key hallmarks of ferroptosis, including lipid peroxidation, macro-blebbing, as well as associated features including ROS production and MMP. The variable responses observed between the two cell lines, even to established ferroptosis inducers, likely reflect differences in their baseline redox state, lipid composition, or iron metabolism, consistent with the well-established context-dependence of ferroptosis ^34^.

### Thermal proteome profiling identifies TXNRD1 as the primary target of Bostrycin

To identify direct protein targets of bostrycin, we performed two-dimensional thermal proteome profiling (2D-TPP) ^35^ to assess proteome-wide changes in protein thermal stability as a function of both temperature and bostrycin concentration. Specifically, we treated MDA-MB-231 cells with five concentrations of bostrycin for one hour, each followed by exposure to nine melting temperatures. Proteome analysis occurred through a TMT-based approach to determine bostrycin-induced protein stabilization (**Figure 4A**; see methods for details). In total, among more than 4000 proteins that were quantified (**Figure 4B; Supplementary Table S4**), the most stabilized protein was TXNRD1, which occurred both in a temperature and concentration-dependent manner (**Figure 4C** and **Supplementary Figure S4A**), thus exhibiting archetypal melting behavior that strongly suggesting direct physical interaction with bostrycin. TXNRD1 (Thioredoxin reductase 1) fulfills a central role in glutathione metabolism and redox control, and has been closely implicated in ferroptosis ^34^.

**Figure 4.**
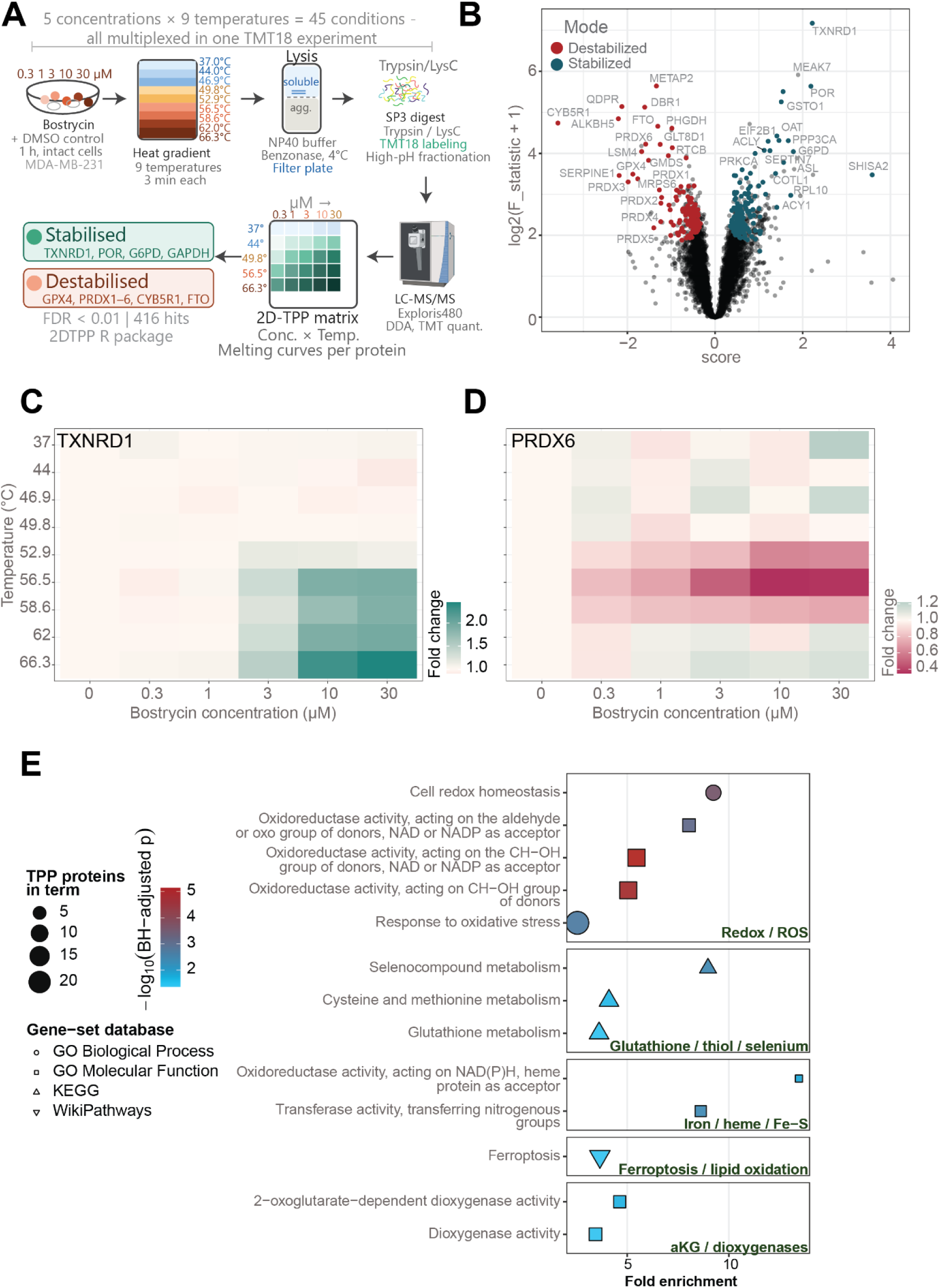
Thermal proteome profiling of bostrycin-treated MDA-MB-231 cells. **A.** Experimental outline. **B.** Volcano plot indicating the results of the 2D-TPP algorithm, with the F-statistic on the y-axis over the score. **C.** Intensity of TXNRD1 as a function of temperature and bostrycin concentration in the TPP analysis. **D.** Same as in D, for PRDX6. **E**. ORA analysis of the TPP experiment.

Beyond TXNRD1, several additional proteins were stabilized or destabilized, suggesting bostrycin-induced alterations in their conformation, interactions, or biochemical state. To gain broader insight into the cellular processes affected, we performed ORA using all 416 TPP hits (FDR <0.01) as input. This revealed pronounced enrichment of processes related to redox regulation, including cellular redox homeostasis, response to oxidative stress, and several oxidoreductase activities (**Figure 4E**). Additional enriched terms involved glutathione, thiol and selenium metabolism, iron/heme-associated functions, ferroptosis, and 2-oxoglutarate-dependent dioxygenase activity. Together, these enrichments indicate that bostrycin-induced thermal shifts converge on interconnected redox-, iron-, and ferroptosis-related processes, supporting widespread perturbation of the cellular redox network.

Several of the most prominently (de)stabilized proteins have been associated with ferroptosis, either directly or indirectly. Strikingly, the destabilized proteins included GPX4, the hallmark negative regulator of ferroptosis, and a selenoprotein like TXNRD1. Other pronounced examples of destabilized proteins included all 6 human peroxiredoxins (PRDX1-6), exhibiting bostrycin concentration-dependent destabilization within a confined temperature range (52.9-58.6°C), as exemplarily shown for PRDX6 (**Figure 4D, and Supplementary Figure S4B**). Peroxiredoxins rely on TXNRD1-mediated reduction via thioredoxin to regenerate their active reduced form. PRDX6 in particular has a recently recognized role in determining ferroptosis sensitivity by interacting with GPX4 ^36^, and by serving as a selenium-acceptor contributing to GPX4 expression ^37^. PRDX destabilization in TPP may be explained by oxidation-driven changes in their multimeric state, as peroxiredoxins exist in a redox-dependent equilibrium between oligomeric forms that may differ in thermal stability (**Supplementary Figure S4C**)^38^. Consistent with this, the thioredoxin-interacting protein TXNIP, which competes with peroxiredoxins for thioredoxin binding and thereby modulates redox homeostasis, was downregulated in the nascent proteome across multiple cell lines (**Figure 2F**), further supporting broad disruption of the TXNRD1-Trx-PRDX axis upon bostrycin treatment (**Supplementary Figure S4D**).

CYB5R1 was among the most strongly destabilized proteins, whereas POR was among the most strongly stabilized (**Figure 4B**). Both are oxidoreductases that promote phospholipid oxidation and thereby induce ferroptosis ^39, 40^. The opposing melting behavior of POR and CYB5R1 may reflect differences in their molecular interactions and resulting conformational states, e.g. POR uses heme as an Fe3+-containing co-factor and NADPH as a substrate, while CYB5R1 uses cytochrome b5 as its Fe3+-containing co-factor and NADH as its substrate. It is striking that several other strongly de-stabilized (but not stabilized) proteins use iron as a co-factor (FTO, ALKBH5, METAP2, PHYHD1, DBR1). Interestingly, they have diverse functions that are distinct from those with a direct redox function, suggesting that their melting behavior may be explained by altered iron availability. In addition, several (de)stabilized proteins depend on alpha-ketoglutarate (αKG) as a substrate (FTO, ALKBH5, PHYHD1, PHGDH (de-stabilized); OAT (stabilized)), a key product of glutaminolysis and metabolic mediator of ferroptosis ^33^. Among these, FTO and ALKBH5 stand out because they both require iron as a co-factor and αKG as a substrate, and because they are the only known m6A RNA demethylases in humans. In addition, both have been firmly implicated in ferroptosis through their control of mRNA methylation and translation of key ferroptotic regulators ^41, 42^. Our data now suggest that the reverse can also occur: bostrycin-induced ferroptosis affects FTO and ALKBH5 stability, potentially through altered iron or αKG availability.

Collectively, our 2D-TPP data indicate that bostrycin induces stability imbalance among multiple proteins in the core of cellular redox regulation, with TXNRD1 as the most likely direct target. The observation that several established ferroptotic drivers and mediators, including GPX4, are among the destabilized proteins suggests that bostrycin induces ferroptosis by acting through engaging the redox metabolic network.

### Metabolome analysis indicates reduced cysteine availability

As our proteome analysis indicated that bostrycin affects several metabolic processes, we performed a targeted metabolomic analysis of MDA-MB-231 and MCF7 cells treated with either DMSO, bostrycin, or IKE for 24 h **(Figure 5A; Supplementary Table S5).** Overall, we identified and quantified 78 polar metabolites, including molecules linked to redox homeostasis and central carbon metabolism.

**Figure 5.**
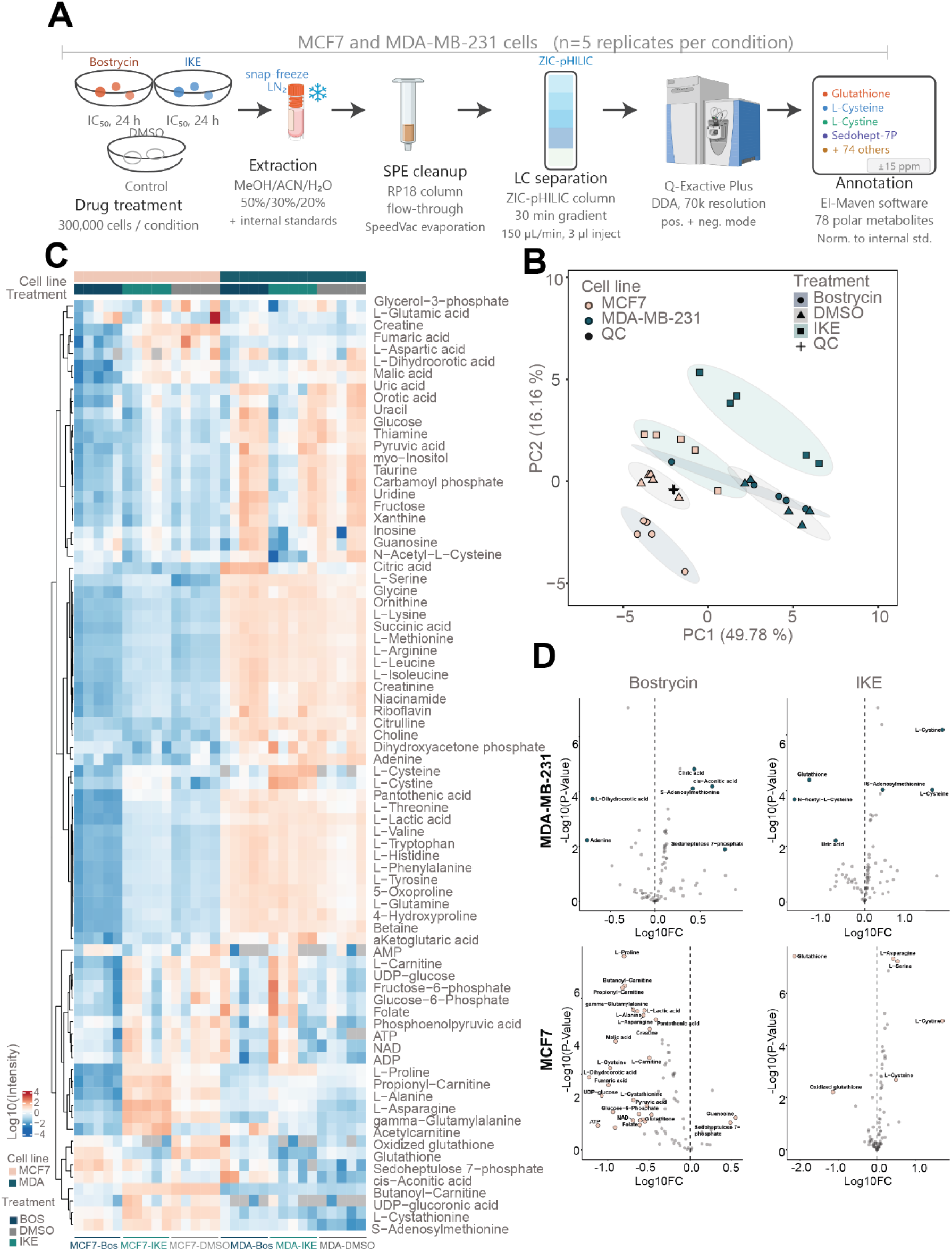
Metabolome analysis of bostrycin-treated MDA-MB-231 and MCF7 cells. **A.** Heatmap showing z-scored metabolite intensities upon treatment of MDA-MB-231 and MCF7 cells with bostrycin or IKE. Row-wise clustering was performed using the pearson-method. **B.** PCA analysis of the metabolome analysis of MDA-MB-231 cells (green) and MCF7 cells (beige) after bostrycin (circle) or IKE (square) treatment (both IC50) for 24h. **C.** Z-scored heatmap of all quantified metabolites in MCF7 cells comparing Bostrycin treatment with DMSO control at IC50 concentration for 24h. Row-wise clustering was performed using the pearson-method. **D.** Volcano plots indicating differential abundance of metabolites after bostrycin (left panels) or IKE treatment (right) in MDA-MB-231 (top) and MCF7 cells (bottom). Highlighted are metabolites that changed in expression more than two-fold and exhibited a p-value < 0.05.

PCA of the metabolomic data revealed treatment-dependent responses that differed substantially between the two cell lines **(Figure 5B).** In MCF7 cells, both bostrycin- and IKE-treated samples separated from their respective DMSO controls. In MDA-MB-231 cells, the effect of bostrycin was more modest and showed substantial overlap with the control, whereas IKE induced a markedly stronger separation. Consistent with these cell-type-dependent responses, hierarchical clustering of metabolite intensities revealed distinct treatment-associated metabolic patterns **(Figure 5C),** with bostrycin eliciting broader metabolic changes in MCF7 cells than in MDA-MB-231 cells.

Differential metabolite analysis further demonstrated distinct responses between treatments and cell lines **(Figure 5D)**. IKE treatment decreased glutathione (GSH) while increasing cystine and cysteine in both MDA-MB-231 and MCF7 cells. Although GSH depletion is consistent with impaired antioxidant capacity during ferroptosis, the increase in cystine and cysteine is unexpected given that IKE inhibits cystine uptake through system xc^−^ and may reflect compensatory redistribution within the metabolic network. Bostrycin induced a comparatively limited number of significant metabolite changes in MDA-MB-231 cells, with a profile clearly distinct from that observed following IKE treatment, supporting different mechanisms of action. In contrast, MCF7 cells exhibited pronounced metabolic remodeling following bostrycin treatment, characterized predominantly by decreased metabolite abundance, including glutathione, cysteine, and several amino acids **(Figure 5D)**.

Cysteine was clearly decreased in MCF7 cells and showed a similar reduction in the metabolite profile of MDA-MB-231 cells, although the effect was less pronounced. Reduced cysteine availability is consistent with a consequence of TXNRD1 inhibition and provides a potential mechanistic connection to both glutathione metabolism and iron-sulfur cluster biogenesis. Cysteine is required for glutathione synthesis and also serves as the sulfur donor for iron-sulfur cluster assembly; consequently, reduced cysteine availability could simultaneously compromise cellular antioxidant capacity and iron homeostasis.

Sedoheptulose 7-phosphate (S7P) increased following bostrycin treatment in both cell lines **(Figure 5D)**. S7P is an intermediate of the pentose phosphate pathway and has previously been reported to accumulate during ferroptosis ^43^. Its accumulation therefore points to perturbation of the pentose phosphate pathway, which provides NADPH required for cellular redox defense.

Collectively, these results demonstrate that bostrycin induces strongly cell-type-dependent metabolic remodeling, while altered cysteine availability emerges as a common and mechanistically relevant feature linking TXNRD1 inhibition to impaired glutathione metabolism and potentially to the disruption of iron-sulfur cluster biogenesis described below.

### Bostrycin induces an iron starvation response

A defining feature of ferroptosis is its dependence on iron. Cellular iron is primarily stored in ferritin complexes, with a small fraction present as a cytosolic labile iron pool (LIP). Because free iron can drive lipid peroxidation via the Fenton reaction, LIP levels are tightly controlled ^44^. Iron homeostasis is maintained by the IRP/IRE system by coordinating iron uptake and storage (**Supplementary Figure 4A**). Under iron-deficient conditions, IRP1 and IRP2 stabilize 3′ IRE-containing transcripts such as TFRC, promoting iron uptake, while repressing translation of 5′ IRE-containing transcripts such as FTH1 and FTL, thereby limiting iron storage ^45^.

Strikingly, exactly this iron-starvation signature was consistently observed in our nascent proteome data across all eight cell lines after bostrycin treatment: proteins encoded by mRNAs with repressive IREs, including ferritin heavy chain (FTH), were downregulated, whereas transferrin receptor (TFRC), encoded by an activating IRE-containing mRNA, was upregulated (**Figure 6A**). This coordinated protein-level response, spanning simultaneous suppression of iron storage and activation of iron uptake, strongly indicates that bostrycin induces an iron-starvation program. Strikingly, treatment of MCF7 cells with the canonical ferroptosis inducers IKE and RSL3 minimally affected FTH and TFRC expression (for IKE) or even resulted in an opposite expression pattern compared to bostrycin (for RSL3) (**Figure 6B**), indicating the absence of an iron starvation response, or even export of iron. These results strongly suggest that bostrycin-mediated iron starvation is a driver, and not a consequence, of ferroptosis.

**Figure 6.**
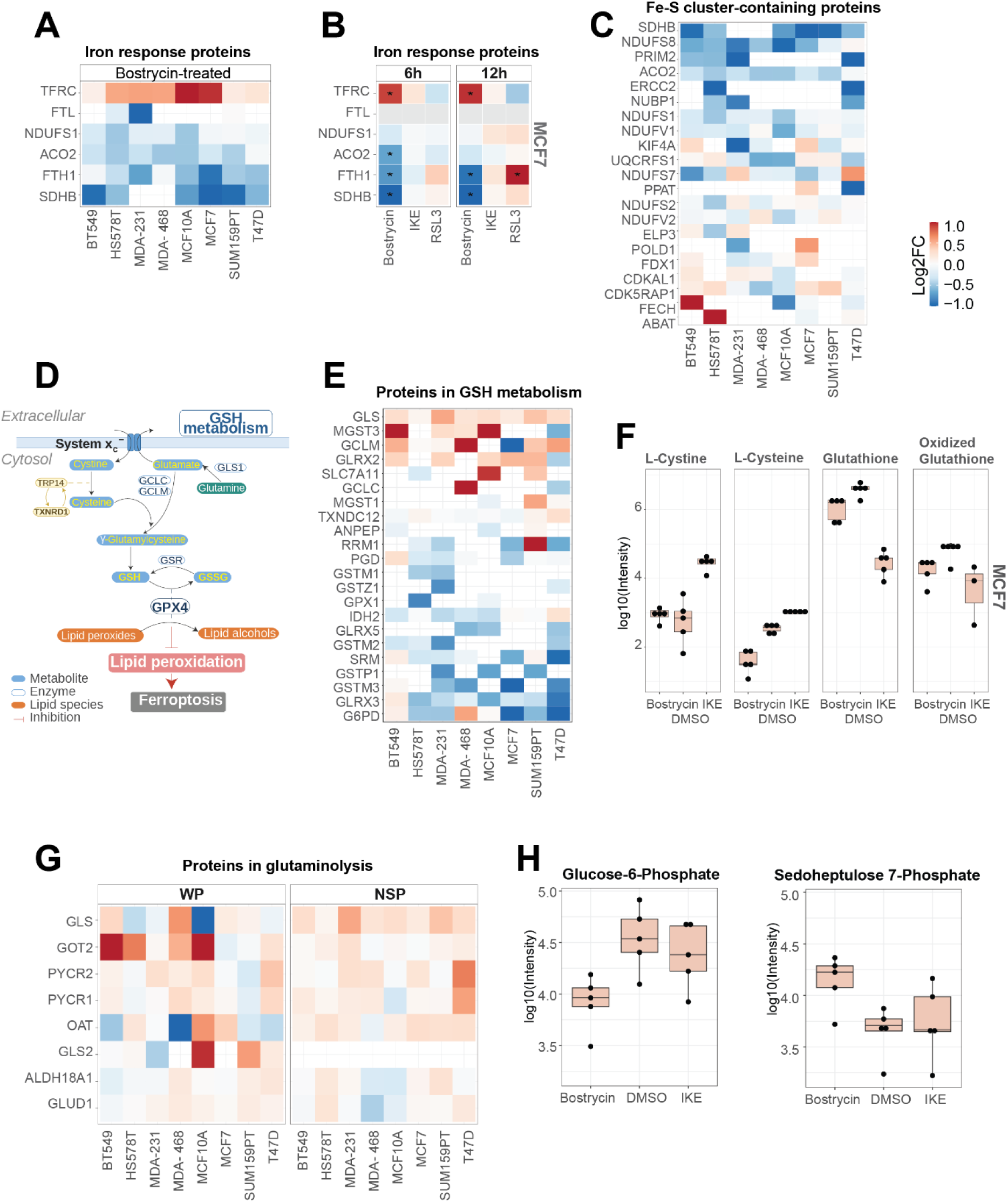
Multi-omics analysis of bostrycin-induced inner defense mechanisms and the regulatory penumbra. **A**. Tile plot of nascent proteomic data upon bostrycin treatment, indicating the average log2 fold change of proteins encoded by IRE-containing mRNA. **B.** Tile plot of nascent proteomic data showing the average log2 fold change of proteins encoded by IRE-containing mRNAs in MCF7 cells treated with bostrycin, IKE or RSL3 for 6 h and 12 h. Grey tiles indicate proteins not quantified in at least two of three replicates. Asterisks mark significantly changed proteins (limma, p < 0.05 and |log2 fold change| > 0.58). **C.** As in A, for iron sulfur-cluster-containing proteins. **D.** Schematic representation of GSH metabolism. **E.** As in A, for proteins involved in the GSH metabolism. **F.** Metabolome analysis of L-Cystine, L-Cysteine, Glutathione and Oxidized Glutathione in MCF7 cells after 24h of bostrycin or IKE treatment at IC50 concentration. **G.** As in A, for proteins involved in glutaminolysis based on either the global proteome (left) or the nascent proteome analysis (right) after treatment with bostrycin at IC50 concentration for 6h (global proteome) and 12h (nascent proteome). **H.** Boxplot indicating the abundance of Glucose-6-phosphate and Sedoheptulose-7-Phosphate after 24h of either bostrycin or IKE treatment at IC50 concentration.

Notably, the observed iron-starvation response contrasts with NRF2-mediated iron regulation, which typically upregulates ferritin to sequester free iron and limit Fenton chemistry ^46^. Instead, our data suggest that bostrycin promotes iron overload through activation of the iron-starvation response rather than NRF2-mediated iron sequestration, thereby sensitizing cells to ferroptosis. Consistent with this interpretation, concurrent TFRC upregulation and FTH downregulation has previously been shown to enhance ferroptosis sensitivity ^47^.

Providing further support for iron accumulation, our nascent proteome analysis showed upregulation of HMOX1 alongside downregulating BLVRA in all eight cell lines (**Figure 2H**), a combination shown to promote iron release during heme catabolism ^48^. The downregulation of BLVRA is particularly notable, as it has recently been identified as a non-enzymatic binding partner and activator of NRF2, such that its depletion attenuates NRF2 signaling ^49^. This provides a mechanistic explanation for why NRF2 target genes involved in iron sequestration remain suppressed despite NRF2 upregulation, further shifting the iron regulatory balance toward iron accumulation.

The iron-starvation response described above may itself be a consequence of disrupted iron-sulfur (Fe-S) cluster biogenesis. Fe-S clusters are essential cofactors for many proteins, particularly in the TCA cycle and electron transport chain (ETC) ^50^. Bostrycin-mediated inhibition of TXNRD1 reduced the availability of cysteine (**Figure 5C, D**), the sulfur donor for Fe-S clusters, suggesting the inhibition of Fe-S cluster formation. Indeed, our nascent proteome data revealed a general decrease in Fe-S cluster–containing proteins upon bostrycin treatment (**Figure 6C**). Impeded Fe-S cluster assembly has been shown to induce an iron starvation response and to sensitize cells to ferroptosis ^51^, which is consistent with our observations (**Figure 6A**). Together, these findings suggest a mechanism wherein bostrycin induces an iron-starvation program by limiting cysteine availability through TXNRD1 inhibition, thereby disrupting Fe-S cluster biogenesis and triggering the iron-starvation response as a downstream consequence **(Supplementary Figure 4A).**

### Bostrycin engages inner defense mechanisms and the regulatory penumbra of ferroptosis

The experimental approaches and data layers presented above—including the global and nascent proteomes, thermal proteome profiling (TPP), metabolomics, and functional assays—each provided distinct evidence supporting the induction of ferroptosis by bostrycin. In particular, this was evidenced by observing that bostrycin elicits macro-blebbing, ROS production, and lipid peroxidation (**Figure 3A-E**). In the model of Dixon and Pratt these properties constitute the core hallmarks of ferroptosis, which receive input from a complex interplay of diverse additional cellular functionalities referred to as inner defense mechanisms and the regulatory penumbra ^52^. We used this model as a framework to integrate our datasets and propose a comprehensive mechanistic understanding of bostrycin-induced ferroptosis.

#### Glutathione metabolism

The GSH axis has been known as the main protector against ferroptosis, targeted by the main inducers of ferroptosis, erastin and RSL3, that either inhibit import of cystine through the system xc^−^ or inhibit lipid peroxidation through GPX4. The synthesis of GSH is a multi-step process that starts with the import of cystine (**Figure 6D**). Upon treatment with bostrycin, most proteins involved in this process were downregulated in our nascent proteomic data, with a few exceptions (**Figure 6E**). The metabolome analysis of MCF7 cells revealed that levels of L-Cysteine, Glutathione (GSH) and oxidized glutathione (GSSG) are downregulated (**Figure 6F**). The concurrent decrease in both GSH and GSSG suggests depletion of the total glutathione pool rather than a shift in redox equilibrium alone. Together, the combination of nascent proteome and metabolome data showed a disruption in the synthesis of GSH upon bostrycin treatment, thereby impairing one of the most important protection layers against ferroptosis.

#### Glutaminolysis

We investigated how bostrycin impacted other metabolic pathways, in particular glutaminolysis, glycolysis and the pentose phosphate pathway (PPP). Import and conversion of glutamine into TCA cycle intermediates is an important driver of ferroptosis, mediated by glutamine uptake, glutaminolysis, and TFRC-mediated transferrin import ^53^. Our global and nascent proteome analyses showed the broad upregulation of glutaminolytic enzymes across all cell lines (**Figure 6G**), along with enhanced TFRC expression as already noted above (**Figure 6A**), consistent with enhanced glutaminolytic flux known to support ferroptosis. Collectively, our data indicate that bostrycin enhanced glutaminolysis.

#### Glycolysis and the pentose phosphate pathway

Our proteomic data also indicated wide-spread changes in expression and synthesis of proteins in glycolysis and the PPP. The first enzymes of glycolysis (HK2, GPI) tended to be increased across cell lines, while those lower in the pathway were decreased (**Supplementary Figure S5B**). In the PPP, enzymes in the oxidative branch (G6PD, H6PD) were increased, while those in the non-oxidative branch (RPIA, RPE, TALDO) showed a general decrease across the tested cell types (**Supplementary Figure 5C)**. Together, this suggests a shift away from glycolysis in favor of the PPP. Interestingly, phosphofructokinase (PFKP) and glucose-6-phosphate 1-dehydrogenase (G6PD), rate-limiting enzymes for glycolysis and the PPP, respectively ^54, 55^, appeared as top-hits in the TPP analysis: G6PD was strongly stabilized upon bostrycin treatment, while PFKP was strongly destabilized (**Figure 4B**). Although it is difficult to infer functional implications, this observation may be explained by altered protein conformation or interactions with substrates as a result of altered pathway activity. The metabolomic data of MCF7 cells showed that bostrycin treatment decreased glucose-6-phosphate (G6P), the substrate of G6PD and the branching point to the PPP, and increased the level of sedoheptulose-7-phosphate (S7P) (**Figure 6H)**, a non-oxidative PPP intermediate. These observations are consistent with enhanced flux through the oxidative PPP, possibly serving to increase NADPH production previously shown as a defense against ferroptosis ^56, 57^. These data support the hypothesis that glycolysis is shifted to the PPP to generate more reducing equivalents in the form of NADPH.

#### TCA cycle and energy metabolism

Since our data indicated that bostrycin affected glutaminolysis and glycolysis as core metabolic pathways that feed into the TCA cycle and energy metabolism, and since bostrycin was previously implicated to interfere with the TCA cycle ^16^, we examined in detail how bostrycin affected individual proteins and metabolites in the TCA cycle and the electron transport chain (ETC). Proteins in the TCA cycle showed moderate upregulation upon bostrycin treatment, with the exception of aconitate hydratase 2 (ACO2) and succinate dehydrogenase B (SDHB), which were strongly and consistently downregulated across the 8 cell lines (**Figure 7**). The fact that ACO2 and SDHB were the only downregulated TCA proteins, and the only TCA enzymes containing IRE sequences, strongly suggests that their downregulation reflects the iron starvation response rather than a general metabolic effect (**Figure 6A**). In addition, both SDHB and ACO2 contain Fe-S clusters, and they thus additionally rely on iron for their enzymatic activity. This is also the case for several other ETC subunits in complex I, II (including SDHB), and III, and strikingly these are all downregulated upon treatment with bostrycin (**Figure 7**). Their downregulation thus suggests compromised Fe-S cluster formation or availability. This contrasts with proteins in complex IV and V, which show consistent upregulation in all eight cell lines (**Figure 7**). This suggests increased activity of the electron transport chain, which is in line with our observation of an increased MMP (**Figure 3F**).

**Figure 7.**
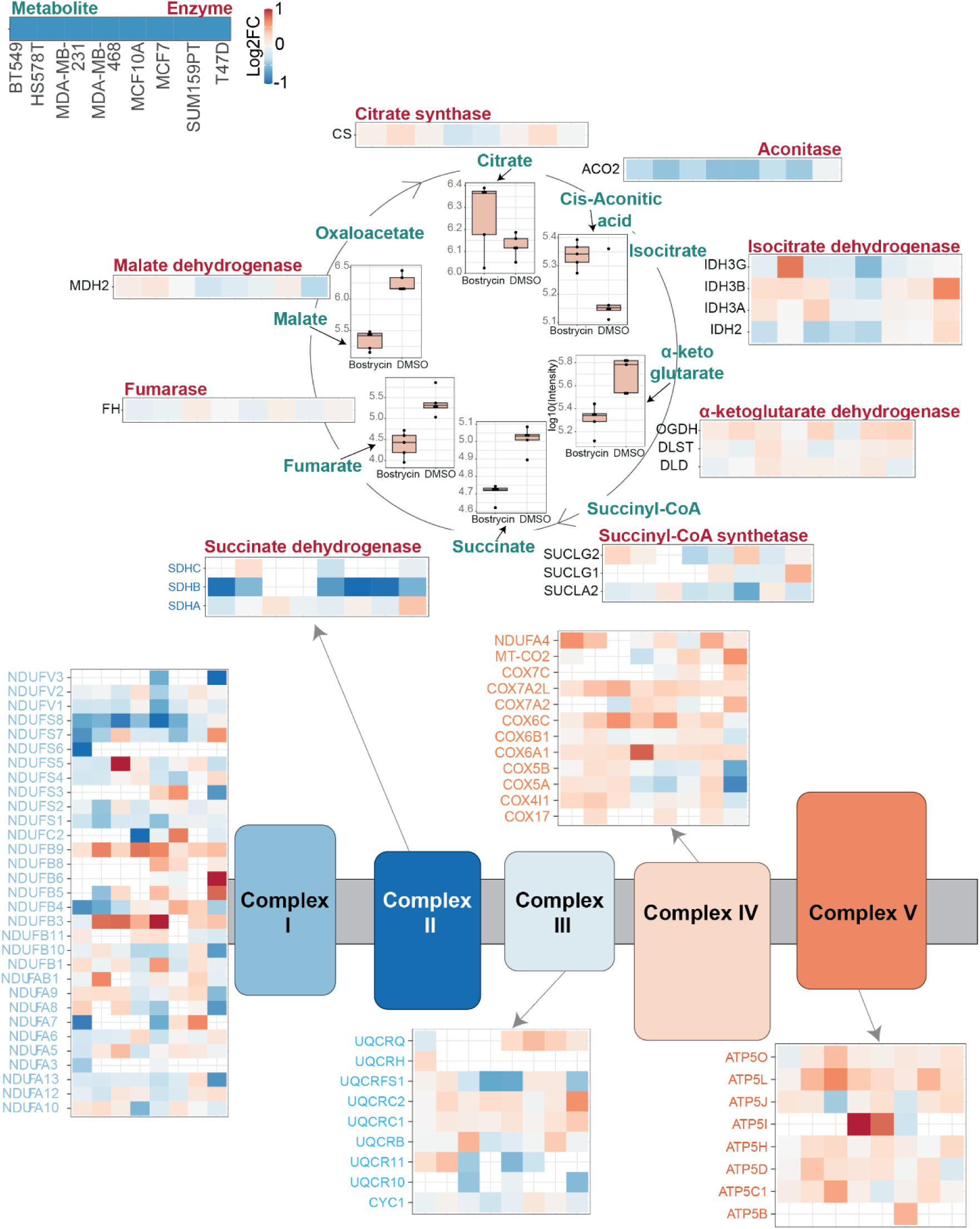
Bostrycin-induced proteome and metabolome alterations in the TCA cycle or the electron transfer chain (ETC). Tile plots indicate the log2 fold change for each protein or protein complex is upon 12h of bostrycin treatment at IC50 concentration for each cell line. For each metabolite, boxplots indicate the changes in metabolite abundance after treatment of MCF7 cells with bostrycin for 24h.

At the metabolomic level, bostrycin led to reduced levels of most TCA metabolites (**Figure 7**), indicating altered availability of TCA intermediates possibly as a consequence of downregulated ACO2 and SDHB. This included aKG, the TCA cycle intermediate at which glutaminolysis-derived carbon enters the cycle. Reduced αKG levels may also explain the thermal destabilization of several αKG-dependent enzymes detected in the TPP analysis (**Figure 4B**). Collectively, our proteomic and metabolomic data indicate how the bostrycin-induced ferroptotic response connects TCA cycle activity and oxidative phosphorylation through iron-dependent gene, protein and metabolite regulation.

### Integrative model

Our TPP data identifying TXNRD1 as the primary target of bostrycin, together with proteomic, metabolomic, and functional evidence for an iron-starvation response and ferroptosis, allow us to propose an integrated model for bostrycin-induced cell death (**Figure 8**). By targeting TXNRD1, bostrycin impairs thioredoxin-mediated redox homeostasis, leading to reduced cysteine availability. This has two interconnected consequences: first, decreased glutathione (GSH) levels increase susceptibility to ferroptosis; second, limited cysteine impairs mitochondrial Fe-S cluster biogenesis, triggering an iron-starvation response that elevates the labile iron pool and promotes ferroptosis. These effects coincide with broader metabolic perturbations that affect the PPP, TCA cycle, and ETC. Collectively, this model demonstrates that bostrycin induces ferroptosis through tightly linked redox, iron, and metabolic pathways, illustrating the power of a multi-omics approach to elucidate complex mechanisms of compound action.

**Figure 8.**
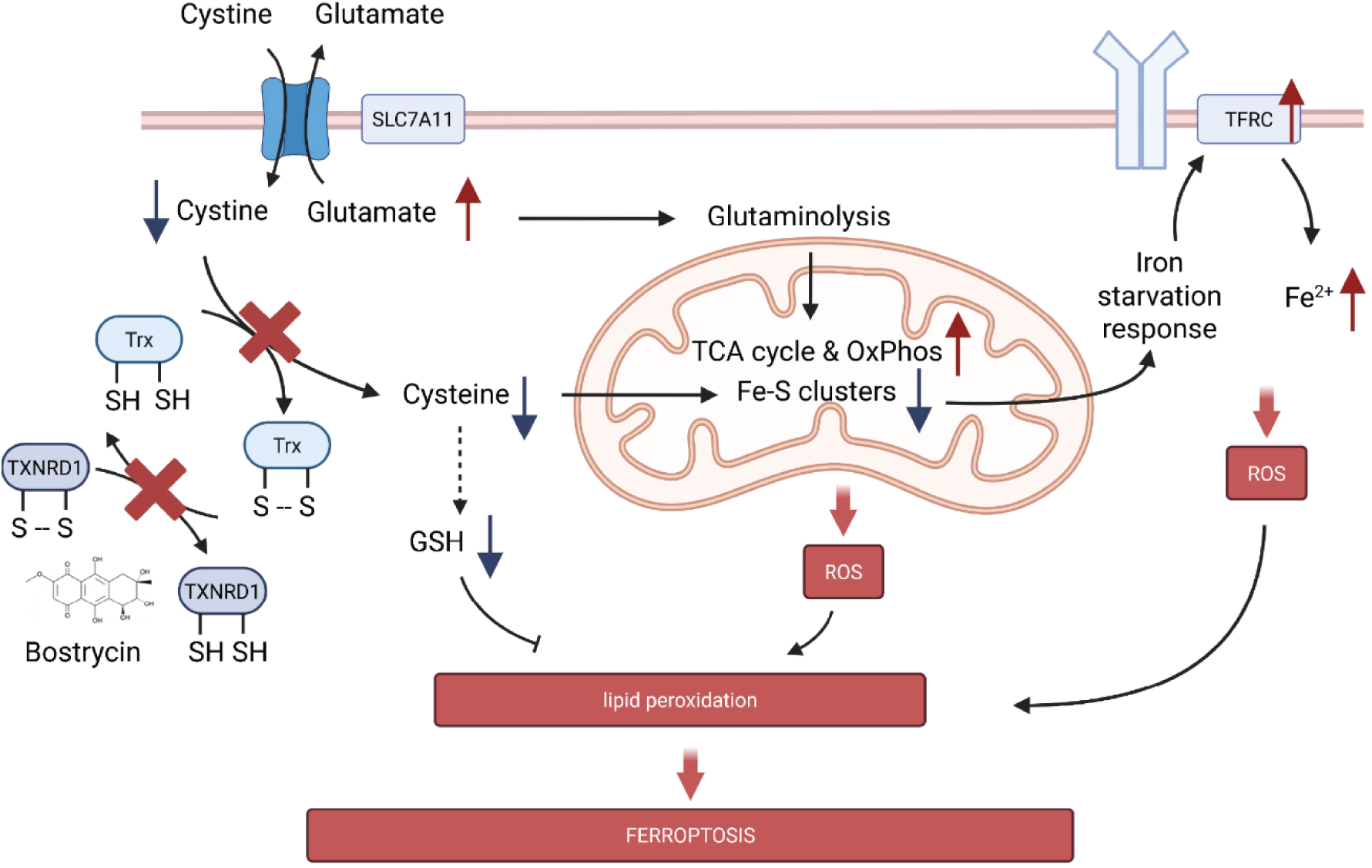
**Integrative model** of proteomic and metabolomic data explaining how bostrycin induces ferroptosis. Created with BioRender.com

## Discussion

The main aim of this study was to understand the mechanism of action by which the small natural compound bostrycin triggers cell death in cancer cells. To approach this broadly, we applied a multi-omics approach integrating global and nascent proteomics, thermal proteome profiling, metabolomics, and functional assays. While each layer on its own hinted that bostrycin induces ferroptosis, only their combination revealed the full scope of cellular pathways underlying the mechanism of action. In particular, proteomic and nascent proteomic data indicated the induction of an oxidative stress response, involving multiple players including those implicated in ferroptosis (**Figure 2G**). Using thermal proteome profiling, a key finding was the identification of bostrycińs primary direct target TXNRD1, an enzyme at the core of the ferroptotic machinery that reduces cystine to cysteine, serving as a precursor for GSH as well as Fe-S clusters. Consistent with TXNRD1 inhibition, reduced cysteine levels were observed in our metabolomic data (**Figure 5C, D**), and the proteomic data revealed a strong iron-starvation response signature (**Figure 6A**). Functional experiments confirmed that bostrycin induces key hallmarks of ferroptosis, including ROS production, lipid peroxidation, and mitochondrial membrane hyperpolarization (**Figure 4**). In addition, our integrated molecular analyses provide deeper insight into the metabolic networks underlying bostrycin activity.

Ferroptosis is canonically induced by intracellular iron overload resulting in membrane lipid peroxidation, however it also receives input from various other environmental and metabolic cues, including redox homeostasis ^52, 58^. Given its role in cysteine and GSH metabolism, bostrycińs primary target TXNRD1 is central to this network and has been previously recognized as a regulator of ferroptosis ^59^. In fact, other compounds have been shown to induce ferroptosis by inhibiting TXNRD1 ^60^, and it has even been suggested that TXNRD1, not GPX4, is the target of RSL3, an established ferroptosis inducer ^61^. Metabolic changes in MCF7 cells stood out by the large number of downregulated metabolites, including cysteine. Limited cysteine availability may be explained by reduced conversion of cystine into cysteine, as observed before upon TXNRD1 inhibition ^62^. Reduced cysteine levels may provide a crucial mechanistic link in our model since it can explain several of the observations in our study, and since it plays a central role in ferroptosis: i) as a precursor of GSH, which is used by GPX4 as a lipid peroxidase, and ii) as a sulfur donor for Fe-S cluster formation in mitochondria, where decreased cysteine levels lead to iron-sulfur cluster deficiency, in turn triggering iron starvation response and ferroptosis ^63, 64^. Specifically, disruption of NFS1, the cysteine desulfurase responsible for extracting sulfur from cysteine for Fe-S cluster biogenesis, as well as its activator frataxin (FXN), have been shown to compromise iron-sulfur cluster formation, and to induce a feedback iron starvation response to increases iron import. This was evidenced by concurrent increased levels of transferrin receptor (TFRC) and decreased levels of ferritin (FTH), in turn elevating ferroptosis sensitivity ^47, 51^ – closely matching our observations. A further causal link between iron starvation response and induction of ferroptosis was confirmed by the observation that deletion of IRPs, the IRE-binding proteins initiating the iron-starvation response, abrogates ferroptosis ^65^. In line with this, indeed several studies have shown that limited availability of cysteine is the trigger of this ferroptotic cascade ^63, 64, 66^. The NRF2 pathway is often induced as a response to redox stress and controls the expression of genes to maintain iron homeostasis and prevent ferroptosis. Despite the increased expression of NFE2L2 (NRF2) (**Figure 2H**), the decreased rather than increased expression of proteins encoded by some of its target genes (most notably FTL, FTH and BLVRA) suggest that NRF2 activity is overridden by alternative mechanisms that result in iron overload instead of sequestration. This is in strong agreement with a recent study showing that inhibition of TXNRD1 induced an NRF2 response but could not prevent cell death ^67^. Key observations in our study show the induction of a strong iron starvation response and dysregulation of Fe-S-cluster-containing proteins mirror these findings, suggesting that bostrycin induces ferroptosis by this mechanism. While TXNRD1 is a target gene of NRF2, the observed stabilization in TPP is unlikely the result of NRF2-signaling given the short exposure (1h) to bostrycin in TPP. Moreover, (de)stabilized proteins in TPP did not include other NRF2 targets observed upon extended exposure (6-12h) in nascent proteome analysis.

The TPP analysis not only revealed TXNRD1 as bostrycińs primary direct target (**Figure 4B, C**) but also showed prominent thermal shifts of several other proteins involved in ferroptosis (e.g. GPX4, CYB5R1, POR) and redox control (e.g. PRDX1-6) (**Figure 4B, E)**. The observed thermal shifts can likely be explained by altered protein conformation resulting from oxidative stress, altered iron availability, or by changes in interactions with substrates or co-factors, and can be generally interpreted as a dis-balanced redox-control network induced by bostrycin (**Figure 4E**). Interestingly, TPP also revealed thermal shifts of proteins with a more distant role to ferroptosis, providing an opportunity to gain insight into additional players and mechanisms beyond those already known to be involved in ferroptosis. In this respect, FTO and ALKBH5, both of which were strongly destabilized in the TPP analysis, may present an intriguing case: these two proteins are the only known human RNA m⁶A demethylases, and for their enzymatic activity they both rely on iron as well as αKG ^68^, i.e. central intermediates in ferroptosis. Moreover, FTO and ALKBH5 have been previously shown to regulate ferroptosis by demethylating, and thereby regulating the transcription, of transcripts encoding key ferroptotic factors including GPX4 ^69, 70^, SLC7A11 ^71, 72^, FSP1 ^73^ and ACSL4 ^74^. This places FTO and ALKBH5 within the emerging framework of epigenetic regulation of ferroptosis mediated by m⁶A methylation ^75^. Beyond the known role of FTO and ALKBH5 in regulating ferroptosis, our thermal proteome data suggest that the reverse relationship may also exist: bostrycin treatment alters iron and αKG availability, and therefore their binding to FTO and ALKBH5. While the mechanistic basis for the FTO thermal shift remains less clear, the thermal stability of ALKBH5 has been shown to increase upon binding to N-Oxalylglycine, an αKG mimetic ^76^. Therefore, the bostrycin-induced thermal shift of ALKBH5 may reflect the loss of its interaction with αKG, potentially altering its demethylase function. Whether ferroptosis-associated changes in iron and αKG availability affect FTO and ALKBH5 activity remains to be demonstrated, but establishing this link would support a regulatory feedback loop between ferroptosis and epigenetic control of ferroptotic gene expression, as previously proposed ^77^. Collectively, our results underscore that TPP is a powerful discovery tool for detecting changes in protein states that reflect perturbation-induced shifts in metabolic processes ^78^, complementing conventional expression-based proteomics.

Although we propose a unified mechanism for bostrycin-induced ferroptosis here, considerable heterogeneity was observed between the eight tested cell lines. This reflects the well-established context-dependence of ferroptosis and suggests that underlying pathways may be wired in a cell type-dependent manner. Conversely, patterns that were consistent across all eight cell lines, such as proteins involved in the iron starvation response (**Figure 6A**) reinforced that this is a core cellular program induced by bostrycin independent of cell-type-specific metabolic context. Importantly, the absence of an iron starvation response signature in MCF7 cells treated with the canonical ferroptosis inducers IKE and RSL3 (**Supplementary Figure S3D**) strongly suggests that bostrycin-mediated iron starvation is a driver, and not a consequence, of ferroptosis. More generally, differences in proteome (**Supplementary Figure S3**) and metabolome response (**Figure 5**) between bostrycin, IKE and RSL3 indicate that there is no unified molecular signature that defines ferroptosis, and that cellular adaptation depends on the mechanism of the ferroptotic inducer. This was also evident in functional assays, where bostrycin induced lipid peroxidation to an extent equal to (in MDA-MB-231 cells) or greater than (in MCF7 cells) that induced by IKE and RSL3 (**Figure 3B, C**). Similarly, cell-type and treatment-dependent differences in ROS production and mitochondrial membrane potential indicate that context-dependent responses extend to processes that are more peripherally related to ferroptosis.

In conclusion, our multi-omics analysis reveals that bostrycin induces ferroptosis through direct targeting of TXNRD1 and activation of an iron-starvation response, via a mechanism distinct from canonical ferroptosis inducers. Our proteomic and metabolomic data provide detailed insight into the metabolic networks intersecting with the core ferroptotic machinery and, more broadly, demonstrate how integrated multi-omics approaches can elucidate the mechanisms of action of uncharacterized natural compounds with widespread metabolic effects.

## Methods

### Cell culture

MCF7, MDA-MB 231, MDA-MB 468, SUM159PT, HS578T, BT549 were cultured in high glucose DMEM (Dulbecco’s Modified Eagle Medium), 10% fetal bovine serum (FBS), 1% penicillin/streptomycin (P/S) and 1x GlutaMAX, while T47D cells were kept in RPMI-1640 media supplemented with 10% FBS, 1% P/S and 1x GlutaMAX. MCF10A cells were cultured in DMEM:F12 (DMEM-Nutrient mixture F-12), 5% horse serum, 20 ng/ml Epidermal Growth Factor, 0.5 mg/ml Hydrocortisone, 100 ng/m Choleratoxin, 10 µg/ml Insulin and 1% P/S. All cells were grown at 37°C with 5% CO2, the medium was exchanged every second to third day and experiments were, , carried out at around 70% confluency if not indicated otherwise. The triple negative breast cancer cell lines MDA-MB 231, MDA-MB 468, SUM159PT, HS578T and BT549 were a kind gift from the lab of Loayza Puch, DKFZ, while MCF7 and MCF10A cells were kindly provided by the Scholl lab, DKFZ. Media components were purchased from LIFE Technologies if not indicated otherwise. Powder for the preparation of SILAC media was purchased from Enzo Life-Sciences and isotopically labeled amino acids bought from Silantes. Drugs were purchased from the following vendors: Bostrycin (Biomol, Cay27466), IKE (Biomol, Cay27088), RSL3 (Hölzel Diagnostika, A15865), Ferrostatin-1 (Biomol, Cay17729) and Liproxstatin-1 (Biomol, Cay17730).

### IC50 determination

To determine the inhibitory concentration of bostrycin after 72h, cell viability was assessed using the CellTiterGlo assay (Promega, G7572). Cells of each cell line were seeded in a 96 well plate with cell densities of 5000 cells/well. After 24 hours, each cell line was subjected to a range of drug concentrations between 0.01 – 128 µM in triplicates. After 72h the cell viability was assessed using the CellTiterGlo assay. For that, media was aspirated and 30 µl of CellTiterGlo solution was added to each well. After 10 min incubation including 2 minutes of shaking at 1200 rpm on an orbital shaker, luminescence was measured using the Mithras LB940 plate reader. IC50 values were calculated using a non-linear regression model using GraphPad Prism software Version 8.0. The option” log(inhibitor) vs. normalized response – Variable slope [Y=Bottom + (Top-Bottom)/(1+10^((LogIC50-X)*HillSlope))]” was chosen.

### Co-treatment assay

9000 MDA-MB-231 cells and 12000 MCF7 cells per well were seeded in a 96 well plate. On the next day, cells were treated in quadruplicates with the indicated concentrations of drugs (Bostrycin: 40 µM, IKE: 20 µM, Fer-1: 20 µM, Lip-1: 10 µM, Z-VAD-FMK 20 µM, Necrosulfonamide 10 µM) and drug combinations for 24 and 6 hours respectively. Afterwards, cell viability was assessed as described above. Intensities were normalized to control cells that received only DMSO treatment.

### Reactive Oxygen Species, lipid peroxidation and Mitochondrial Membrane Potential determination

30000 or 150 000 MDA-MB-231 or MCF7 cells, respectively, were seeded in a 12-well plate. On the next day, cells were treated with Bostrycin, IKE, RSL3 or DMSO as control for the indicated times and concentrations (see Table 1). CM-H2DCFDA (Invitrogen, C6827), Bodipy C11 (Invitrogen, D3861) or TMRE (Tetra-methylrhodamin-Ethylester, Selleck Chemicals GmbH, E2978) were added to a final concentration of 5 µM, 2 µM or 200 nM, respectively, for further 30 minutes of incubation. After trypsinization, cells were washed, resuspended in PBS containing 1% FBS and kept in the dark and on ice until measurements using flow cytometry-based analysis on the BD FACSCanto II™ (BD Biosciences). For each condition, 10000 single cells were analyzed. At first, living cells were isolated using the side- and forward scatter. In the next step, forward scatter height over area was used to determine single cells and finally the experimental outcome was assessed by analysis of fluorescence intensity captured through the BL-530/30 filter. Gates were set according to the respective controls. For data analysis, FloJo (version 10, BD) was used.

### Global proteome analysis

For analyzing the global proteome of all cell lines upon bostrycin treatment, cells were seeded in 10-cm plates and on the next day, treated with media containing their respective IC50 concentrations of bostrycin for 6 hours. After harvesting, cells were lysed in 0.1% Rapigest in 100 mM ammonium bicarbonate with 10 mM CAA and 40 mM TCEP. After sonication (Benson probe sonicator, 18%, 5 s on, 7 s off, 2 min on time), a BCA assay was conducted with the cleared lysate according to manufacturer’s instructions (Pierce™ BCA Protein Assay Kits, Thermo Scientific, 23225). Trypsin/LysC was added to 100 µg of protein and incubated overnight. The digestion was stopped using a final concentration of 1% TFA. After further 30 min incubation at 37°C, samples were cleared through centrifugation for 30 min at 16 000g. Peptides were then subjected to mass spectrometry analysis.

### Nascent proteome analysis

For the nascent proteome analysis, cells were seeded in 10 cm plates in triplicates for each condition. On the next day, if pre-treatment is indicated, cells were pre-treated with the drug at IC50 value or DMSO as control for 6 hours. Afterwards cells were deprived of Arginine (Arg), Lysine (Lys) and Methionine (Met) by incubating with Arg, Lys and Met-free media (Enzo Life Sciences, AES-0430, AES-0432 or AES-0433) for 45 minutes to ensure depletion of intracellular stores. Then, cells were treated with heavy or intermediate Arg/Lys-containing media (Silantes, 211104113, 201204102, 201604102 or 211604102) together with AHA (Jena Bioscience, CLK-AA005-500) instead of Met, and either the drug at IC50 concentration or DMSO as control for additional 6 hours. In case of no pre-treatment, the protocol was carried out starting with the amino acid deprivation step. After the indicated treatment times, cells were harvested by scratching, cells were pelleted by centrifugation and flash-frozen liquid nitrogen.

Sample preparation was conducted using the Click-It alkyne-agarose enrichment kit (LifeTechnologies, C10416). Samples were first lysed in CHAPS-containing urea lysis buffer (200 mM HEPES pH8, 0.5 M NaCl, 4% CHAPS, 8 M Urea) and sonicated on ice using a Benson probe sonicator (10 sec on/10 sec off time, 2 minutes total on time, 10% pulse amplitude). Lysates were cleared by a centrifugation step (30 min at 15 000g and 4°C). After determining the protein concentration using a Pierce™ BCA Protein Assay according to manufacturer’s instructions, 750 µg of each treated and control sample were mixed. The mixed lysates were then alkylated by addition of iodoacetamide for 30 minutes at room temperature and afterwards, 100 µl of alkyne-agarose slurry and 93 µl of the reaction mixture for the [3+2] cyclo addition (10 µl Cu(II)SO4 (200 mM), 62.5 µl tris (hydroxypropyltriazolylmethyl)amine (THPTA, 160 mM), 10 µl aminoguanidine (2 M), 10 µl sodium ascorbate (2 M)) were added and incubated at 40°C for 2h. After the click-reaction, the resin was centrifuged at room temperature and all supernatant was discarded, since all newly synthesized proteins are covalently bound to the alkyne beads. Beads were then briefly washed with water and 2 ml of reduction-alkylation solution (10 mM Tris(2-carboxyethyl)phosphine (TCEP), 40 mM 2-Chloroacetamide (CAA) in SDS wash buffer) was added and incubated for 15 minutes first at 70°C and for additional 15 minutes at room temperature. After reduction and alkylation, the beads are thoroughly washed. In total, five iterations of 1ml SDS-Wash buffer (100 mM Tris-HCl pH 8.0, 1% SDS, 250 mM NaCl, 5 mM EDTA) were followed by 1 ml of water, five iterations of 1 ml Guanidine-Wash buffer(100 mM Tris-HCl pH 8.0, 6M Guanidine-HCl) and additional five iterations of Acetonitrile-Wash buffer (20% Acetonitrile, ULCMS in Water). The cleaned beads were resuspended in 200 µl digestion buffer (100 mM Tris-HCl pH 8, 2 mM CaCl2, 5% Acetonitrile) and 1 µg Trypsin/LysC mix (Promega, V5073) was added. Proteins were then digested overnight in a shaking incubator for 16h at 37°C. On the next day, the peptide-containing supernatant was collected and acidified using 10% Formic acid to a pH of 2-3. The peptides were then loaded on a prepared Oasis PRiME HLB µElution Plate (Waters, 186008052). The plate was conditioned by addition of 200 µl Acetonitrile, followed by 3 washing steps, first with 200 µl 60% Methanol, 1% formic acid in water and then twice with 1% Formic acid in water. After conditioning, samples were loaded and washed with consecutively applying 750 µl, 250 µl and 100 µl 1% formic acid in water. To elute the peptides, 50 µl of 60% methanol with 1% formic acid was applied followed by 50 µl 100% methanol. The peptides were then dried using the SpeedVac (Savant SPD111V SpeedVac Concentrator, Thermo Scientific) and resuspended in 0.1% formic acid before being measured on the mass spectrometer.

### Thermal proteome profiling

The 2-dimensional (i.e. concentration and temperature-dependent) thermal proteome profiling was carried out essentially as previously described (40). Three million MDA-MB-231 cells were seeded in 10 cm plates and on the next day cells were treated with 0.3, 1, 3, 10 and 30 µM bostrycin or DMSO as control for 1 hour. Cells were first washed with PBS and then harvested by trypsinization and counted. Equal amounts of cells were distributed in a PCR plate, and each set of samples was exposed to one of the following temperatures: 37.0, 44.0, 46.9, 49.8, 52.9, 56.5, 58.6, 62.0 or 66.3 for 3 minutes before placing the cells back on ice. Afterwards, cells were lysed using a benzonase-containing buffer (PBS, 1.33% NP40, 2.5mM MgCl2, 1.6x cOmplete, 1.6x PhosStop, 0.417U/ul benzonase) for 1 hour at 4°C. Aggregated protein was separated from soluble protein using a filter plate (Milipore, MSHVN4550). Soluble proteins were cleaned up and digested using the SP3 protocol.

40 µl of SP3 binding solution (30 µl ethanol, 10 µl 15% formic acid and 2 µl cleaned 1:1 Sera-Mag A and Sera-Mag B bead mixture (Fisher Scientific, 24152105050250 and 44152105050250) was added to a 96 well filter plate and soluble protein samples were added. The mixture was shaken for 15 min at room temperature for complete protein binding. The unbound fraction was discarded by centrifugation (1 min at 1000g). Beads were then washed with 200 µl 70% ethanol four times. For protein digestion, the beads were resuspended in 40µl digestion buffer (100mM HEPES, 0.2 µg LysC, 0.2 µg Trypsin, 5mM CAA and 1.25 mM TCEP) and incubated at RT overnight shaking. On the next day, peptides were eluted from the filter plate and lyophilized for 1 hour. Peptides were then resuspended in water and TMT reagent from a TMT18 kit was added and incubated for 1 hour at room temperature. Non-reacted TMT-reagent was quenched by adding the stop solution (5% Hydroxylamine in water) for 15 min. Samples were then pooled according to the respective experiment. In a last step, peptides were cleaned using the OASIS protocol (see above) and fractionated using the Pierce™ High pH Reversed-Phase Peptide Fractionation Kit (Life Technologies, 84868).

### Mass spectrometry measurements

Measurement of peptides from nascent and global proteome experiments were carried out on an EASY-nLC 1200 system (Thermo Fischer Scientific) coupled to a QExactive HF mass spectrometer (Thermo Fischer Scientific), while the thermal proteome profiling experiment was measured on an UltiMate3000 (Thermo Fischer Scientific) coupled to an Exploris480 (Thermo Fischer Scientific).

When measuring on the QExactive-HF instrument, peptides were loaded on an Acclaim Pepmap C18 5 µm-particle-size trap column (0.1 × 20 mm, Thermo Scientific #164564-CMD) and separated over a 55°C heated nanoEase M/Z Peptide BEH C18 1.7 µm-particle-size analytical column (0.075 × 250 mm, Waters #186008795) with a gradient consisting of buffer A (0.1% FA in H2O) and buffer B (80% ACN in H2O). The gradient employed for nascent proteome approach was as following: The total elution gradient time was 105 min with buffer A (0.1% formic acid in water) and buffer B (0.1% formic acid in 80% acetonitrile). The linear gradient started at 3%B, increasing over four minutes to 8% B, followed by an increase to 10%B in the next two minutes. During minutes 6-74, the gradient rises to 32%B, which further increases to 50%B between minutes 74-86. Within one minute, the gradient then reaches 100% and stays at 100%B for 7 minutes, before dropping back to 3%B for another 10 minutes.

The global proteome was measured on a 210-minute gradient with the following steps: After three minutes of gradient at 2%B, the concentration of solvent B linearly increased to 25% for 150 minutes. During the next 30 minutes solvent B concentration rose to 40% only to reach 95% B within one minute. At the end, five minutes at 95% B were followed by 20 minutes at 2%B. The flow rate for all measurements was set to 300 nl/min.

Separated peptides were injected by electrospray ionization and analyzed by MS/MS with an ion transfer tube temperature set to 275°C and spray voltage set to 2.5 kV. The nascent proteome was measured in data-dependent analysis mode with the following settings: MS1 was measured with a resolution of 60 000 within the range of 375-1500 m/z, with an AGC target of 3000000 and a maximum injection time of 50 ms. The top 20 precursor peaks were selected for MS2 analysis with an AGC target of 100000, a maximum injection time of 50 ms and a resolution of 15000 full width-half maximum.

Global proteome analysis was carried out in data-independent analysis mode. MS1 measurement was performed at a resolution of 60000, with a scan range of 400-1200 m/z and an AGC target of 3000000, while having a maximum injection time of 20 ms. Targeted MS/MS spectra were acquired at a resolution of 30000 while using 34 non-overlapping isolation windows of 24.3 m/z. Fixed mass was set to 200 m/z and the normalized collision energy to 27.

The Thermal Proteome Profiling was measured on a Exploris480 (Thermo Fischer Scientific) coupled to an UltiMate 3000 RSLC nano-HPLC (Dionex). Solvent A and B were prepared as above 0.1% FA in H2O and 80% ACN in H2O, respectively and peptides were separated on a Waters BEH column (75 µm× 250 mm, C18, 1.7 µm, 130 Å ). A 120 min gradient with a flow rate of 300 nl/min was used. In the beginning of the gradient, solvent B was increased from 2% to 6% for 1 min, then to 10% for 6 min and increased linearly over 85 minutes to 25%. Between minutes 95 and 106 solvent B concentration further increased to 42%B, before reaching 80%B for 3 minutes followed by 5 minutes at 2%B. Data was acquired in a data-dependent mode with a fixed cycle time of 2s. MS1 scans were carried out at a resolution of 120000 over a scan range of 375-1500 m/z. The maximum injection time was set to 45 ms, while AGC Target was set at 300% and the RF Lens value was at 40%. MS2 scans were performed at a resolution of 45000 with a fixed first mass of 110 m/z, a maximum injection time of 96 ms, a normalized AGC target of 200% and a RF Lens value of 50%. Normalized HCD collision energy was set to 36% and isolation window was at 0.7 m/z.

### MS data analysis

Raw files from all nascent proteome experiments were searched in MaxQuant version 2.0.1.0 ^79^ using a UniProt human proteome database (reviewed entries only). To account for the SILAC labels, multiplicity was set to 3 with a light channel, an intermediate channel with Arg6 and Lys4, as well as a heavy channel with Arg10 and Lys8. Additionally, Match-between runs and Requantify were enabled. Otherwise, all settings were used at default. To infer proteins and peptides for all global proteome analyses, the search engine DIA-NN (version 1.8) was used ^80^. The same human protein database was used together with a contaminant fasta file ^81^ to create a spectral library. Settings were used at their default values.

The analysis of TPP raw data was carried out in MSFragger v13 in Fragpipe version 18 against the human proteome downloaded from Uniprot ^82^. The following search parameters were included: tryptic digestion with maximum 2 missed cleavages, peptide length 7-50, MS/MS tolerance 10ppm, peptide tolerance 20 ppm, fixed modifications: carbamidomethyl on cysteine and TMT18plex on lysine, variable modifications included acetylation on N termini of proteins, oxidation of methionine and TMT18plex on N termini of proteins. Peptide spectrum matches were validated by philosopher version 4.1.0. The false discovery rate was fixed at 1%. The Quantification of TMT was carried out by TMT-Integrator using the default settings.

### Statistical analysis and data representation

For the analysis of all proteome experiments, the software environment R (version 4.3.1) was used with custom scripts. From the search engines, for DDA experiments the “ProteinGroups.txt” file was loaded and for DIA experiments “report.pg matrix.tsv”. For all SILAC experiments, the normalized H/M ratio was further processed, while the DIA report file already reported LFQ-normalized intensity values. Identified contaminants were filtered out. For differential protein expression, proteins with two valid values per condition were taken and subjected to limma analysis ^83^. Differentially expressed proteins that adhered to the cutoff (p-Value <0.05 and |log2 fold change| > 0.58) were submitted to over-representation analysis using the clusterProfiler package or the reactomePA package with the enrichplot package for visualization ^84, 85, 86^. Gene Set Enrichment Analysis was carried out taking the sorted limma results as a ranked list to the GSEA function of the clusterProfiler package ^84^. Overrepresentation analyses and Gene Set Enrichment Analyses were carried out with the data bases from the msigdbr and org.Hs.eg.db packages ^87^. UpSet plots were generated with the UpSetR package version 1.4.0 ^88^ and venn diagrams using the ggvenn package version 0.1.10 ^89^. Heatmaps were generated using the pheatmap package version 1.0.12 ^90^. Thermal Proteome Profiling analysis was carried out using the 2DTPP package ^91^. Statistical analysis for all other experiments was carried out in GraphPad Prism (version 8.4.3) and conducted as indicated in the Result section.

### Metabolome analysis

For the analysis of polar metabolites, 3.0 × 10^5^ MCF7 or MDA-MB-231 cells were seeded. On the next day they were treated with bostrycin or IKE at their respective IC50 concentrations for 24h. Cells were then snap-frozen in liquid nitrogen. For harvest, cells were taken up in a Methanol, Acetonitrile and water (50%/30%/20%) solution containing internal standards (Table 2). Cells were then lysed by sonication, centrifuged and supernatant was transferred onto an equilibrated RP18 SPE column (Phenomenex Strata C18-E, 55 µm, 50 mg / 1 ml, 8B-S001-DAK). The cell pellet was sonicated again and supernatant transferred onto the SPE column. The flow-through from both transitions was collected and the solvents were evaporated using a SpeedVac overnight at 15 °C. Directly before LC-MS measurement, dried metabolites were resuspended in 450 µl of 75% Acetonitrile.

Water-soluble metabolites were analysed by LC-MS/MS using a Q-Exactive Plus quadrupole-Orbitrap mass spectrometer (Thermo) interfaced with an Ultimate 3000 LC system (Thermo). The analytical gradient was carried out as previously described ^92^ using a SeQuant ZIC-pHILIC column (5 µm, 2.1 mm x 150 mm, Millipore Sigma) with solvent A (20 mM ammonium carbonate + 0.1% ammonium hydroxide in water) and solvent B (acetonitrile). 3 µl of each sample was injected and the analytical gradient lasted 30 minutes, as follows: linear gradient from 80% to 20% within 20 minutes before returning to 80% in 0.5 min and appended with a 7.5-minute equilibration step. The flow rate was maintained at 150 µL/min. The eluents were analyzed by MS in ESI positive/negative mode with data dependent MS2. The full scan at 70000 resolution (69-1000 m/z scan range, × 106 AGC-Target, 50 ms maximum injection time (maxIT)) was followed by a data dependent MS2 scan at 17.5k resolution (1 × 105 AGC target, 20 ms max IT, 1 loop count, 2 s to 10 s apex trigger, 2 × 103 minimum AGC target, 20 s dynamic exclusion). The HESI source parameters were set as 40 sheath gas flow rate, 15 auxiliary gas flow rate, 1 sweep gas flow rate, spray voltage: 3.0 kV in positive mode, 3.1 kV in negative mode, 320°C capillary temperature, and the heater temperature of auxiliary gas was 120°C. The annotation of the metabolites was performed using the EI-Maven software (Elucidata, https://www.elucidata.io/el-maven) with an offset of ± 15 ppm. The intensity of each target was normalized to the internal standards (positive/negative standards for metabolites) and protein concentration.

## Supplementary Figure legends

**Supplementary Figure S1.**
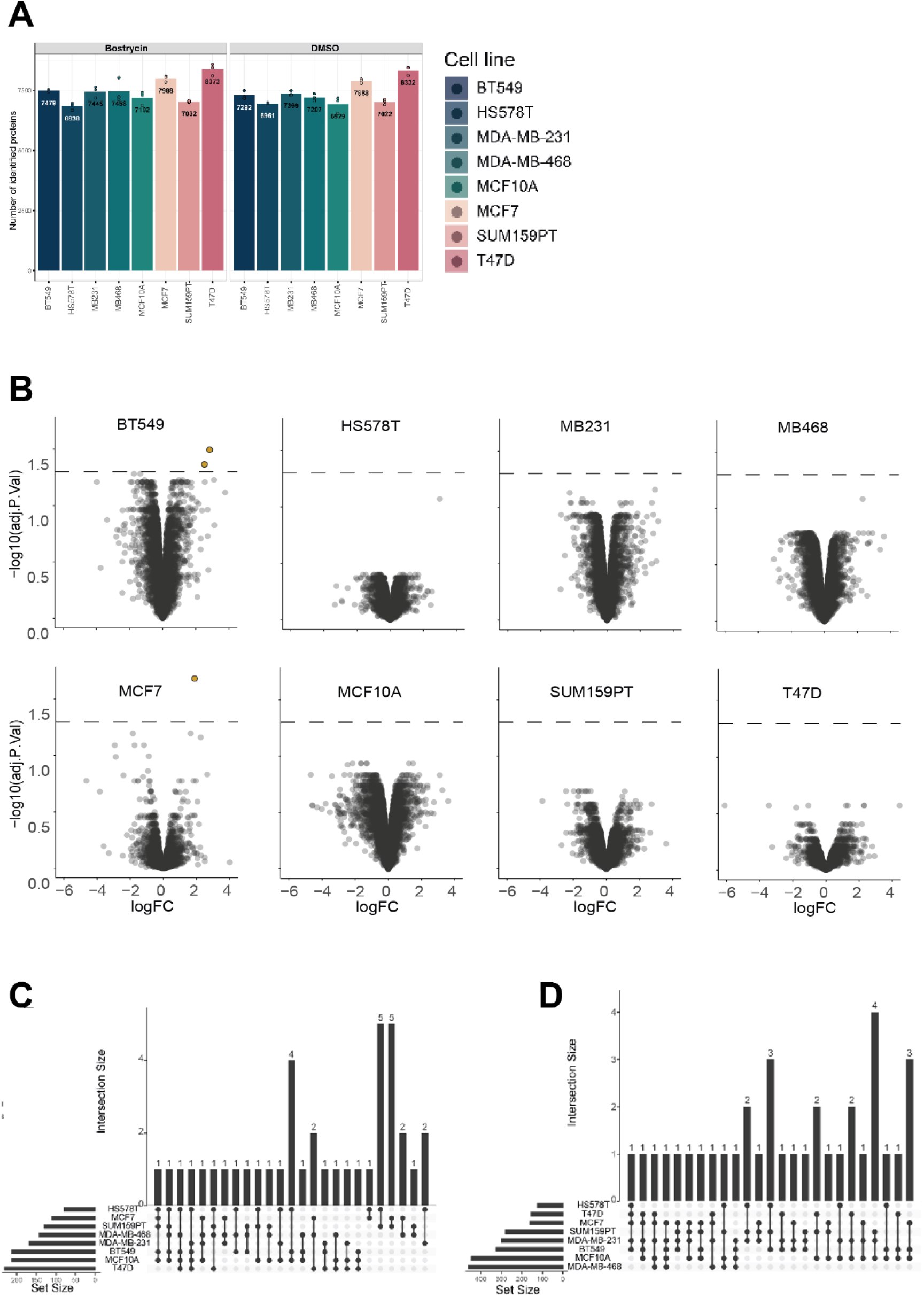
Proteome analysis of bostrycin-treated cancer cells. **A.** Bar plot indicating the number of identified proteins in eight cell lines after treatment with bostrycin (left) or DMSO (right). Dots indicate the number of proteins in independent experiments. **B.** Volcano plots after differential protein expression in bostrycin-treated cancer cell lines. **C, D.** Upset plots indicating the overlap in upregulated (C) and downregulated proteins (D) in bostrycin-treated cancer cell lines.

**Supplementary Figure S2.**
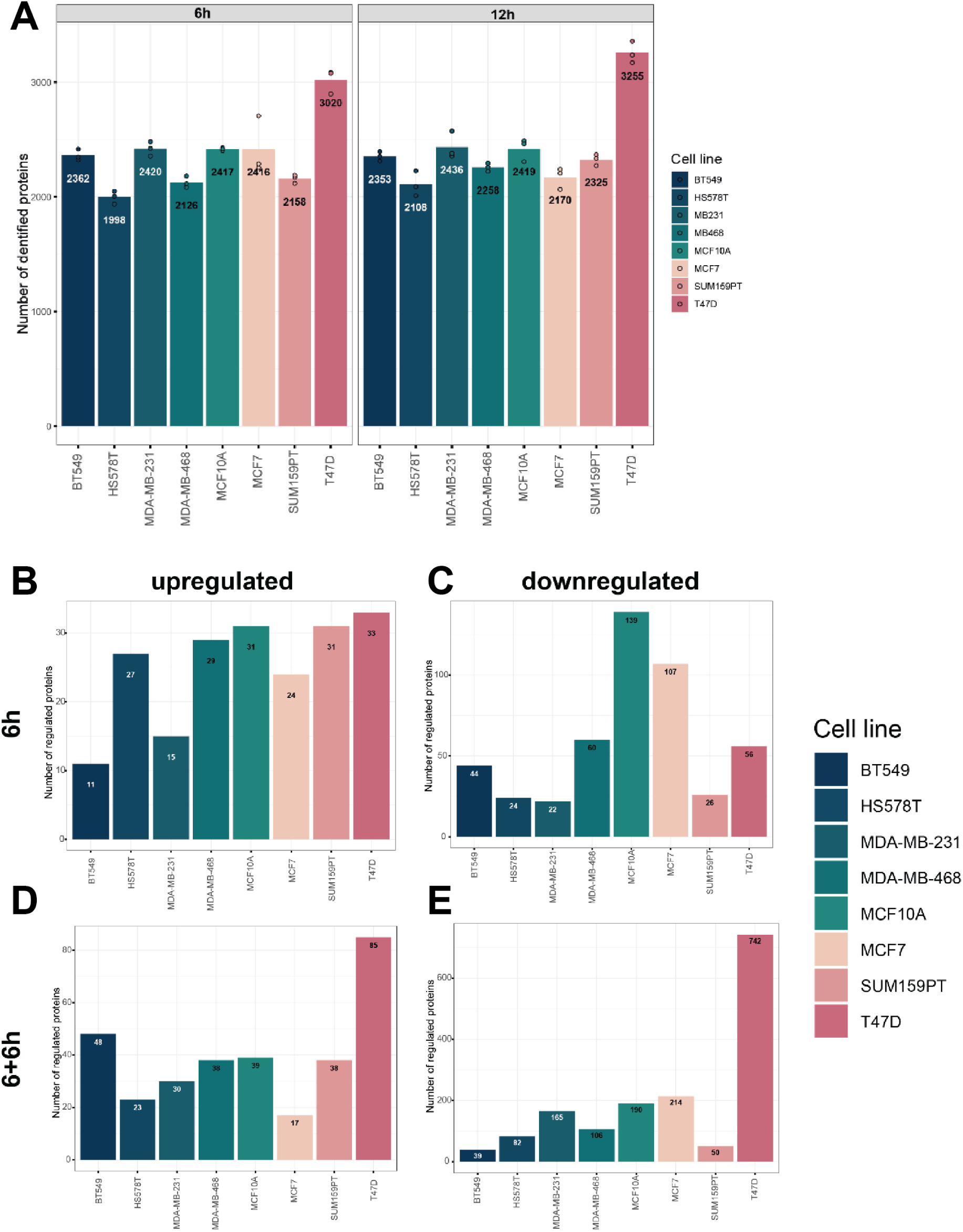
Nascent proteomics of bostrycin-treated cancer cell lines. **A.** Bar plot indicating the number of identified newly synthesized proteins in eight cell lines after treatment with bostrycin for 6h (left) or 12h (right). Dots indicate the number of proteins in independent experiments. **B-D.** Bar plots indicating the number of up (B+D) and down-regulated proteins (C+E) in the nascent proteome after 6h (B+C) or 12h (D+E) treatment with bostrycin treatment. Shown is the number of proteins that increased or decreased in abundance by more than 50% and exhibit a p-value < 0.05 in each cell line and timepoint.

**Supplementary Figure S3.**
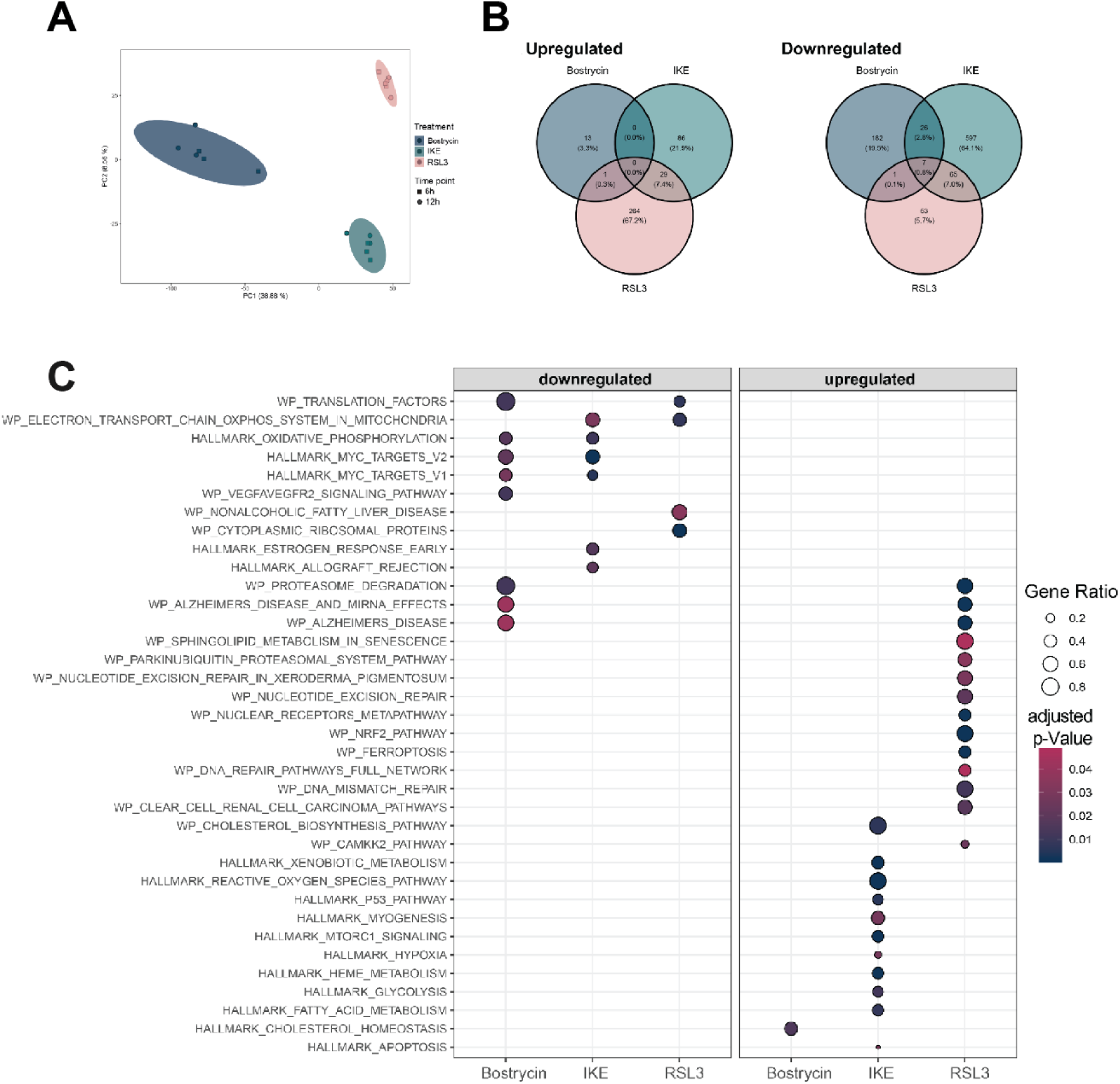
Nascent proteomics after induction of ferroptosis in MCF7 cells. **A.** Principal component analysis of nascent proteomic data from MCF7 cells treated with IKE, RSL3 and bostrycin. **B.** Overlap of up- and downregulated proteins identified in A. **C.** Gene set enrichment analysis of nascent proteomics data acquired after treatment of MCF7 cells with IKE, RSL3 and bostrycin.

**Supplementary Figure S4.**
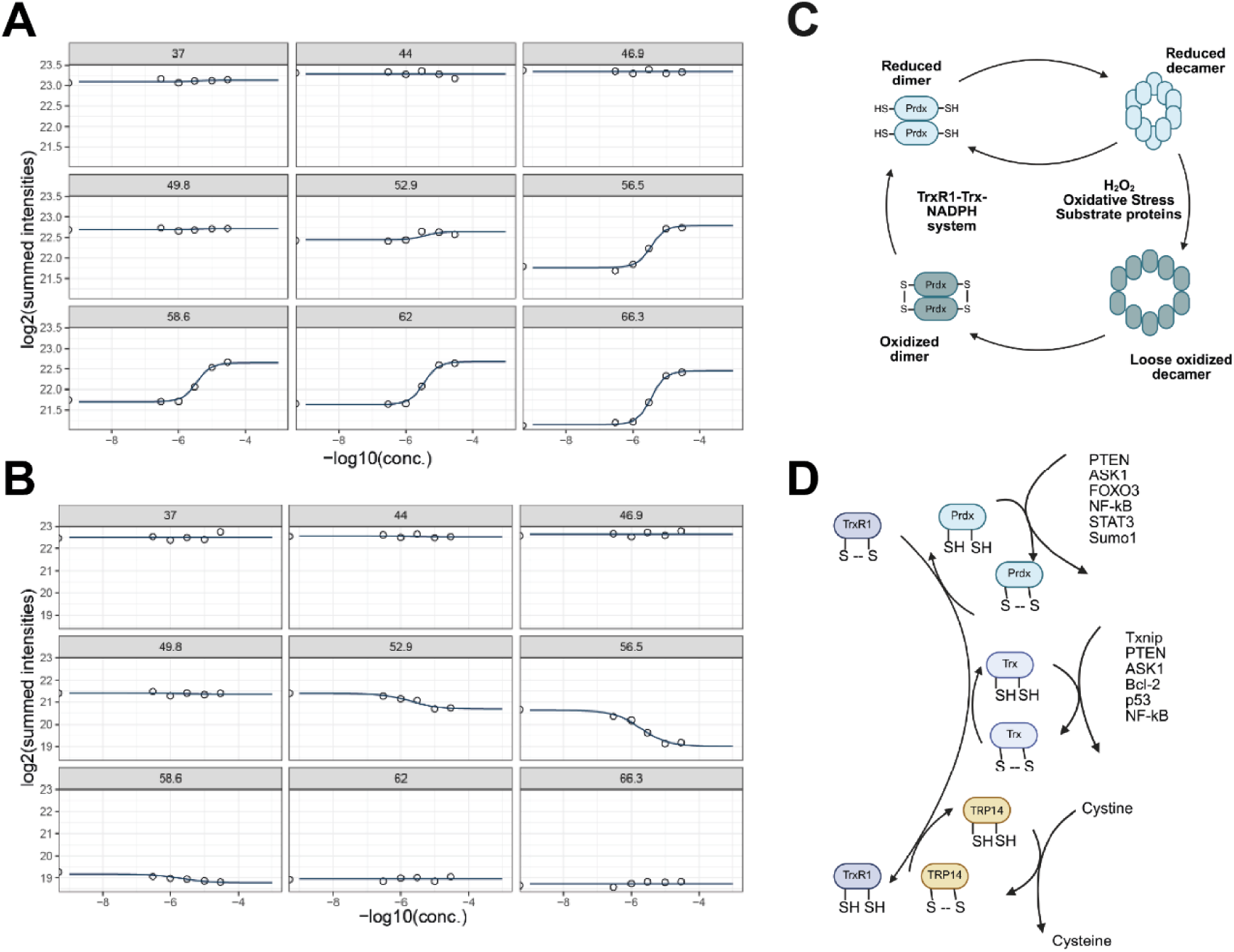
Thermal proteome profiling of bostrycin-treated MDA-MB-231 cells. **A.** Changes in abundance of TXNRD1 in the TPP analysis for each temperature across the different bosytrcin concentrations. **B.** As in A, for PRDX6. **D.** Redox cycle of peroxiredoxins indicating their conformational changes upon oxidation and reduction. **E.** Schematic depiction of TrxR1 and its substrates Prdx, Trx and TRP14. Indicated as well is a selection of their respective binding substrates. D+E created with BioRender.

**Supplementary Figure S5.**
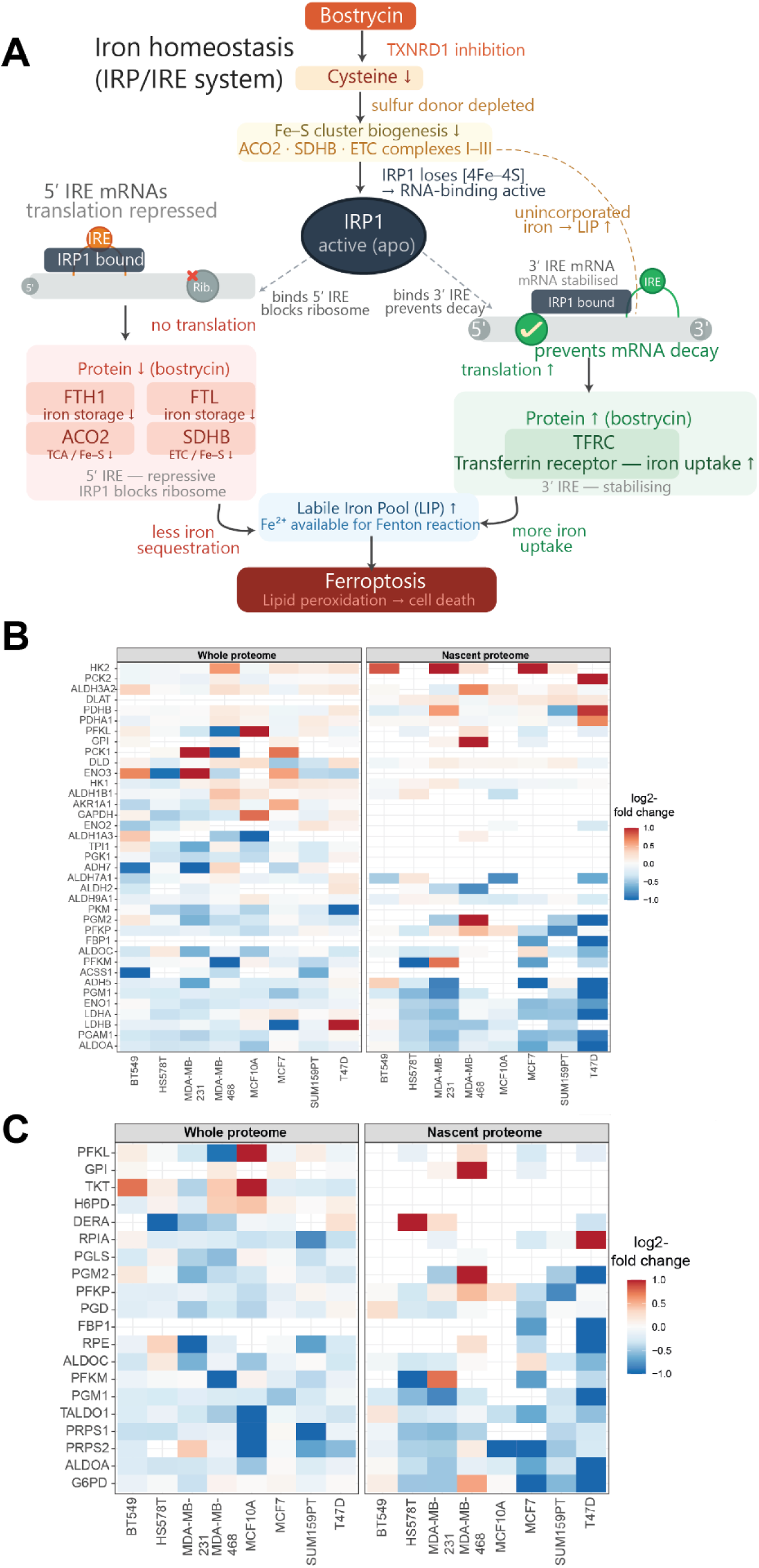
Bostrycin-induced alterations in cell metabolism. **A.** Scheme of iron homeostasis by the IRP/IRE system, and schematic model for bostrycin-mediated iron regulation. **B.** Tile plot of nascent proteomic data upon bostrycin treatment, indicating the average log2 fold change of proteins under the term KEGG_GLYCOLYSIS_GLUCONEOGENESIS. **C.** As in B, for proteins under the term KEGG_PENTOSE_PHOSPHATE_PATHWAY.

## Data availability

Processed global proteome, nascent proteome, thermal proteome profiling and metabolomics data are provided in Supplementary Tables S1–S5. Raw MS data have been deposited with the ProteomeXchange Consortium via the PRIDE partner repository ^93^ with the dataset identifier PXD077097.

## Acknowledgements

J.K. acknowledges funding of this work by the Deutsche Krebshilfe (project 70114190), and by the German Federal Ministry of Research, Technology and Space (BMFTR) as part of the National Research Initiative Mass Spectrometry in Systems Medicine (MSCoreSys, FKZ 03LW0233K). A.S.A. acknowledges support from the Joachim Herz Stiftung through the Add-on Fellowship for Interdisciplinary Science and Transfer. A.S. and J.H. acknowledge funding from the BMFTR as part of SPP2306 (SCHU-2670/2 and SCHU-2670/4) and the German Ministry of Science and Education (BMBF). A.B.C-F was funded by a fellowship from Fundação de Amparo à Pesquisa do Estado de São Paulo (FAPESP 2021/10153-5).

## Author contributions

Conceptualization: L.-M.W., A.S.A., and J.K.; formal analysis: L.-M.W. and A.S.A.; funding acquisition: J.K.; investigation: L.-M.W., A.S.A., J.H., A.B.C.F. and I.B.; methodology: L.-M.W., A.B.C.F. and I.B.; resources: M.S. and A.S.; supervision: M.S., A.S. and J.K.; writing – original draft: L.-M.W. and J.K.; writing – review and editing: A.S.A., I.B., M.S. and A.S., with input from all authors.

## Competing interest

The authors declare no competing interests.

## Ethics declaration

Not applicable.

## Clinical Trial Number

Not applicable.

## Declaration of AI-assisted technologies in manuscript preparation

During the preparation of this work, the authors used ChatGPT to check and refine the English grammar and sentence structure of the manuscript. The authors reviewed and edited the output as needed and take full responsibility for the content of the published article.

## Notes

### Competing Interest Statement

The authors have declared no competing interest.

